# A suberized exodermis is required for tomato drought tolerance

**DOI:** 10.1101/2022.10.10.511665

**Authors:** Alex Cantó-Pastor, Kaisa Kajala, Lidor Shaar-Moshe, Concepción Manzano, Prakash Timilsena, Damien De Bellis, Sharon Gray, Julia Holbein, He Yang, Sana Mohammad, Niba Nirmal, Kiran Suresh, Robertas Ursache, G. Alex Mason, Mona Gouran, Donnelly A. West, Alexander T. Borowsky, Kenneth A. Shackel, Neelima Sinha, Julia Bailey-Serres, Niko Geldner, Song Li, Rochus Benni Franke, Siobhan M. Brady

**Affiliations:** Department of Plant Biology and Genome Center, University of California, Davis, Davis CA 95616, USA; Plant-Environment Signaling, Institute of Environmental Biology, Utrecht University, 3584 Utrecht, the Netherlands; School of Plant and Environmental Sciences, Virginia Tech, Blacksburg, VA 24061, USA; Department of Plant Molecular Biology, University of Lausanne, 1015 Lausanne, Switzerland; Institute of Cellular and Molecular Botany, Rheinische Friedrich-Wilhelms-University of Bonn, Kirschallee 1, 53115 Bonn, Germany; Center for Plant Cell Biology, Department of Botany and Plant Sciences, University of California, Riverside, Riverside, CA 92521, USA; Electron Microscopy Facility, University of Lausanne,1015 Lausanne, Switzerland; Department of Plant Sciences, University of California, Davis, Davis CA 95616, USA; Department of Plant Biology, University of California, Davis, Davis CA 95616, USA

## Abstract

Plant roots integrate environmental signals and developmental programs using exquisite spatiotemporal control. This is apparent in the deposition of suberin, an apoplastic diffusion barrier, which regulates the entry and exit of water, solutes and gases, and is environmentally plastic. Suberin is considered a hallmark of endodermal differentiation, but we find that it is absent in the tomato endodermis during normal development. Instead, suberin is present in the exodermis, a cell type that is absent in the model organism *Arabidopsis thaliana*. Here, we uncover genes driving exodermal suberization and describe its effects on drought responses in tomato, unravelling the similarities and differences with the paradigmatic Arabidopsis endodermis. Cellular resolution imaging, gene expression, and mutant analyses reveal loss of this program from the endodermis, and its co-option in the exodermis. Functional genetic analyses of the tomato MYB92 transcription factor and ASFT enzyme demonstrate the importance of exodermal suberin for a plant water-deficit response. Controlling the degree of exodermal suberization could be a new strategy for breeding climate-resilient plants.

## INTRODUCTION

Plants have evolved complex cell type-specific regulatory processes to respond and adapt to dynamic environments. In certain cell types, such processes allow the formation of constitutive and inducible apoplastic diffusion barriers that regulate mineral, nutrient, and water transport, pathogen entry, and have the capacity to alleviate water-deficit stress (Baxter et al., 2009; Thomas et al., 2007). The *Arabidopsis thaliana* root endodermis contains both lignified and suberized diffusion barriers, of which the latter is extremely responsive to nutrient deficiency (Barberon et al., 2016). Many of the molecular players associated with suberin biosynthesis and the transcriptional regulation of this biosynthetic process have been elucidated using the Arabidopsis root endodermis as a model.

Suberin is a complex hydrophobic biopolymer, composed of a phenylpropanoid-derived aromatic (primarily ferulic acid) and aliphatic (poly-acylglycerol) constituents which is deposited between the primary cell wall and the plasma membrane as a lamellar structure (Molina et al., 2009; Serra and Geldner, 2022). While the order of the enzymatic reactions that produce suberin is not entirely understood (Serra and Geldner, 2022), many of the enzymes associated with suberin biosynthesis have been identified to function in the Arabidopsis root endodermis. These include the elongation of fatty acid acyl-CoA thioesters to very long chain fatty acid-CoA products by a fatty acid elongase complex for which the KCS (ketoacyl-CoA synthase) enzyme DAISY (docosanoic acid synthase) is the rate limiting step (Franke et al., 2009). Suberin primary alcohols are formed by the FAR enzymes (fatty acyl reductases) which reduce C18:0-C22:0 fatty acids to primary fatty alcohols (Domergue et al., 2010). Suberin ω-hydroxyacids (ω-OH acids) and α,ω-dicarboxylic acids are produced by cytochrome P450 monooxygenases from the subfamilies CYP86A, CYP86B and CYP94B which ω-hydroxylate fatty acids (Compagnon et al., 2009; Höfer et al., 2008; Molina et al., 2009). Glycerol esterification of fatty acid acyl CoA derivatives is catalyzed by the glycerol phosphate acyltransferases (GPAT) including GPAT5 (Beisson et al., 2007). Ferulic acid is esterified to ω-hydroxyacids and primary alcohols by the feruloyl transferase ALIPHATIC SUBERIN FERULOYL TRANSFERASE (ASFT)/ω-HYDROXYACID/FATTY ALCOHOL HYDROXY-CINNAMOYL TRANSFERASE (FHT) (Gou et al., 2009; Molina et al., 2009; Serra et al., 2010).

Many of the suberin biosynthetic enzymes were identified based on their co-expression, leading to the hypothesis that a simple transcriptional module coordinates their transcription (Compagnon et al., 2009; Molina et al., 2009; Shukla et al., 2021). Although the overexpression of several transcription factors can drive suberin biosynthesis in either Arabidopsis leaves or roots (Cohen et al., 2020; Kosma et al., 2014), the transcription of suberin biosynthetic genes is redundantly determined. It is only when a set of four Arabidopsis transcription factors are mutated - *MYB41, MYB53, MYB92* and *MYB93* - that suberin is largely absent from the Arabidopsis root (Shukla et al., 2021). Although not studied in roots, the Arabidopsis *MYB107* and *MYB9* transcription factors are required for suberin biosynthetic gene expression and suberin deposition in seeds (Gou et al., 2017; Lashbrooke et al., 2016). These data demonstrate that, in Arabidopsis, multiple transcription factors coordinate the expression of suberin biosynthesis genes, dependent on the organ. Furthermore, these transcriptional regulatory modules are likely conserved across plant species, as orthologs of many of these transcription factors and their target genes are strongly co-expressed across multiple angiosperms (Lashbrooke et al., 2016; Molina et al., 2009).

While the Arabidopsis root endodermis is well characterized anatomically and molecularly, an additional root cell type deposits an apoplastic diffusion barrier during primary growth in other species (Enstone et al. 2002). This cell layer is found below the epidermis, it is the outermost cortical cell layer of the root and has been referred to as either the hypodermis or the exodermis. The latter term was used given observations of a potential Casparian Strip. Indeed, in 93% of angiosperms studied this layer was reported to possess either a lignified Casparian Strip, a suberized diffusion barrier or both (Perumalla et al., 1990). Given the nature of these features, the exodermis is hypothesized to function similarly to the endodermis (Barberon, 2017; Geldner, 2013). The *Solanum lycopersicum* (tomato) root contains both an exodermis and an endodermis. At its first stage of differentiation, a lignified cap is deposited on the outmost (epidermal) face of exodermal cell walls as well as on its anticlinal walls. During its second stage of differentiation, suberin is deposited around the entire surface of the exodermal cells (Kajala et al., 2021). Strikingly, the tomato root endodermis did not contain suberin under the conditions examined. The transcriptional regulators and biosynthetic enzymes associated with this aspect of exodermal differentiation are not known, nor is the influence of root exodermal suberization on environmental stress responses.

Here, we profiled the transcriptional landscape of the tomato exodermis at cellular resolution, characterized suberin accumulation in response to plant hormone abscisic acid (ABA) and in response to water-deficit. We identified a co-expression module of potential suberin-related genes, including transcriptional regulators, and validated these candidates by generating multiple CRISPR-Cas9 mutated tomato hairy root lines using *Rhizobium rhizogenes* (Ron et al., 2014) and tomato plants stably transformed with *Agrobacterium tumefaciens*, and screened them for suberin phenotypes using histochemical techniques. The validated genes included a MYB transcription factor (*SlMYB92, Solyc05g051550*) whose mutant has a reduction in exodermal suberin and the *SlASFT* enzyme, whose mutant has a disrupted exodermis suberin lamellar structure with a concomitant reduction in root suberin levels. To test the hypothesis that suberin is associated with tomato’s drought response, we exposed *slmyb92* and *slasft* mutant lines to water deficit conditions. Both mutants displayed a disrupted response including perturbed stem water potential and leaf water status. This work describes for the first time a genetic mechanism required for exodermal suberin biosynthesis content and integrity, and link these to a plant’s response to reduced water availability.

## RESULTS

### Developmental timing and chemical composition of the tomato suberized exodermis

We previously quantified exodermis suberin deposition along the longitudinal axis of the tomato root (cv. M82, LA3475), using the histochemical stain Fluorol Yellow (FY) In Arabidopsis roots, suberin is absent from the root meristem and elongation zones and begins to be deposited in a patchy manner in the late differentiation zone after the CS has become established, and is then followed by complete suberization in the distal differentiation zone. Quantification of exodermal suberin in seven day old roots demonstrated the same three categories of deposition (none, patchy and complete) (**Figure 1A,C,D; Supplemental Figure 1**). Electron microscopy further demonstrated that within the completely suberized zone, suberin lamellae are deposited primarily on the epidermal and inter-exodermal faces of the exodermal cell (**Figure 1B; Supplemental Figure 1**). Suberin was consistently absent within the root endodermis throughout all developmental zones (**Figure 1A, Supplemental Figure 1**) (Kajala et al., 2021).

**Figure 1:**
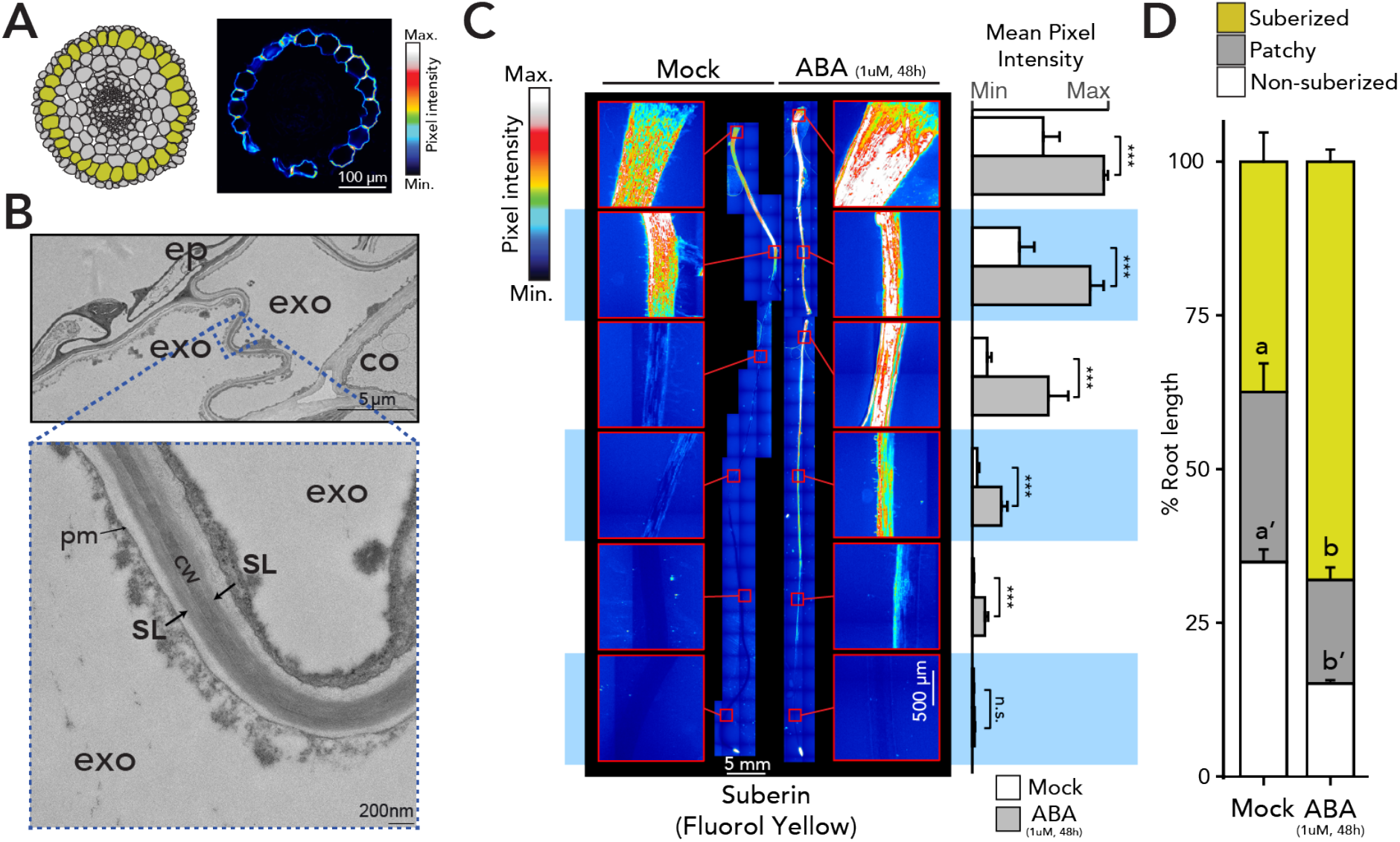
Suberin is deposited in the tomato exodermis and is regulated by ABA. (A) Graphical representation of *S. lycopersicum* (cv. M82) root anatomy (the exodermis is highlighted in yellow) and representative cross section of a 7-day-old root stained with fluorol yellow (FY). Scale bar = 100 μM. (B) Transmission electron microscopy cross-sections of 7-day-old roots obtained at 1 mm from the root-hypocotyl junction. Top image shows the epidermal (ep), exodermal (exo) and inner cortex (co) layers. Bottom image is a close-up of the featured region (zone defined with blue dashed lines), showing the presence of suberin lamellae (SL). cw = cell wall, pm = plasma membrane. (C) Fluorol yellow staining for suberin in wild-type 7-day-old plants treated with mock or 1 μM ABA for 48 h. Whole-mount staining of primary root (left) and quantification of suberin abundance along the root (right), n ≥ 6, error bars: SD. *** = p-value < 0.005. One-way ANOVA followed by TukeyHSD. (D) Developmental stages of suberin deposition of wild-type plants treated with mock or 1 μM ABA for 48 h. Zones were classified as non-suberized (white), patchy suberized (gray) and continuously suberized (yellow), n ≥ 6, error bars: SD.

Monomer profiling of cell wall-associated, and polymer-linked aliphatic suberin monomers in one-month-old tomato roots revealed a predominance of ɑ,ω-dicarboxylic acids, similar to potato (Serra et al. 2010). Compared to Arabidopsis roots, which mostly features ω-OH acids and a maximum chain length of 24 carbons (C24) (Franke et al., 2005, 2009), additional C26 and C28 ω-OH acids and primary alcohols were observed in tomato (**Supplementary Figure 2A**). This phenomenon of inter-specific variation in suberin composition has been previously observed (Kolattukudy et al., 1975). Nevertheless, tomato suberin composition greatly overlaps with Arabidopsis, which should facilitate the identification of orthologous suberin biosynthesis genes.

### Identification of suberin biosynthetic enzymes and transcriptional regulators

In order to map the tomato root suberin biosynthetic pathway and its transcriptional regulators, we leveraged prior observations of relative conservation of transcriptional co-regulation of suberin pathway across angiosperms (Lashbrooke et al. 2016; Molina et al. 2009; Compagnon et al. 2009; Legay et al. 2016). In the Arabidopsis root, suberin levels increase upon treatment with ABA (Barberon et al., 2016; Shukla et al., 2021), a hormone which is a first responder upon water deficit stress. Exodermal suberin deposition in tomato is similarly increased upon ABA treatment, both in terms of the region that is completely suberized as well as in the intensity of the signal (**Figure 1C-D**). *S. lycopersicum*’s wild relative, *Solanum pennellii* (LA0716) is drought tolerant (Eshed et al., 1992; Gong et al., 2010; Gur et al., 2011), and enhanced suberin deposition in Arabidopsis via mutation of *ENHANCED SUBERIN1 (ESB1)* confers drought tolerance, although *esb1* also shows enhanced endodermal lignin and interrupted CS formation (Baxter et al., 2009). Hence, we tested and confirmed the hypotheses that *S. pennellii* has higher suberin deposition than M82 even in water sufficient conditions and shows no changes in suberin deposition pattern in response to ABA in *S. pennellii* seedlings (**Supplementary Figures 2–3**). *S. pennellii* suberin levels are thus constitutive. Therefore, we utilized a gene expression dataset profiling transcription in M82 roots as well as across roots from 76 tomato introgression lines (ILs) derived from *S. lycopersicum* cv. M82 and *S. pennellii* (LA0716) with M82 as the recurrent parent (Toal et al., 2018). We additionally profiled the root transcriptomes of one-month-old tomato plants under well-watered, waterlogged and water-deficit conditions. We hypothesized that genes directly involved in the biosynthesis and deposition of suberin will be highly correlated in both water deficit and the IL population.

By combining both IL, waterlogging, and water deficit datasets in a weighted gene correlation network analysis (WGCNA) (Langfelder and Horvath, 2008), we identified modules of co-expressed genes (**Supplementary Figure 4**). A module (“royalblue”) containing 180 genes was significantly enriched in suberin-related genes (odds ratio = 16.1; p < 0.001). This was confirmed by intersection with the dataset profiling expression in a known activator of tomato fruit suberin biosynthesis (**Supplemental Table 1**) (Lashbrooke et al., 2016). The “royalblue” module contains several orthologs of well-known suberin biosynthetic gene families such as glycerol-3-phosphate acyltransferases (GPATs), 3-ketoacyl-CoA synthases (KCSs), and feruloyl transferases (ASFT/FHTs) (**Figure 2A, Supplemental Table 1**). Additionally, putative tomato orthologs of known transcriptional regulators of suberin biosynthesis, *AtMYB41, AtMYB63* and *AtMYB92 (SlMYB41: Solyc02g079280; SlMYB63: Solyc10g005550; SlMYB92: Solyc05g051550)*, among others, were found in this module (Kosma et al. 2014; Shukla et al. 2021; Cohen et al. 2020; Du et al. 2015).

**Figure 2.**
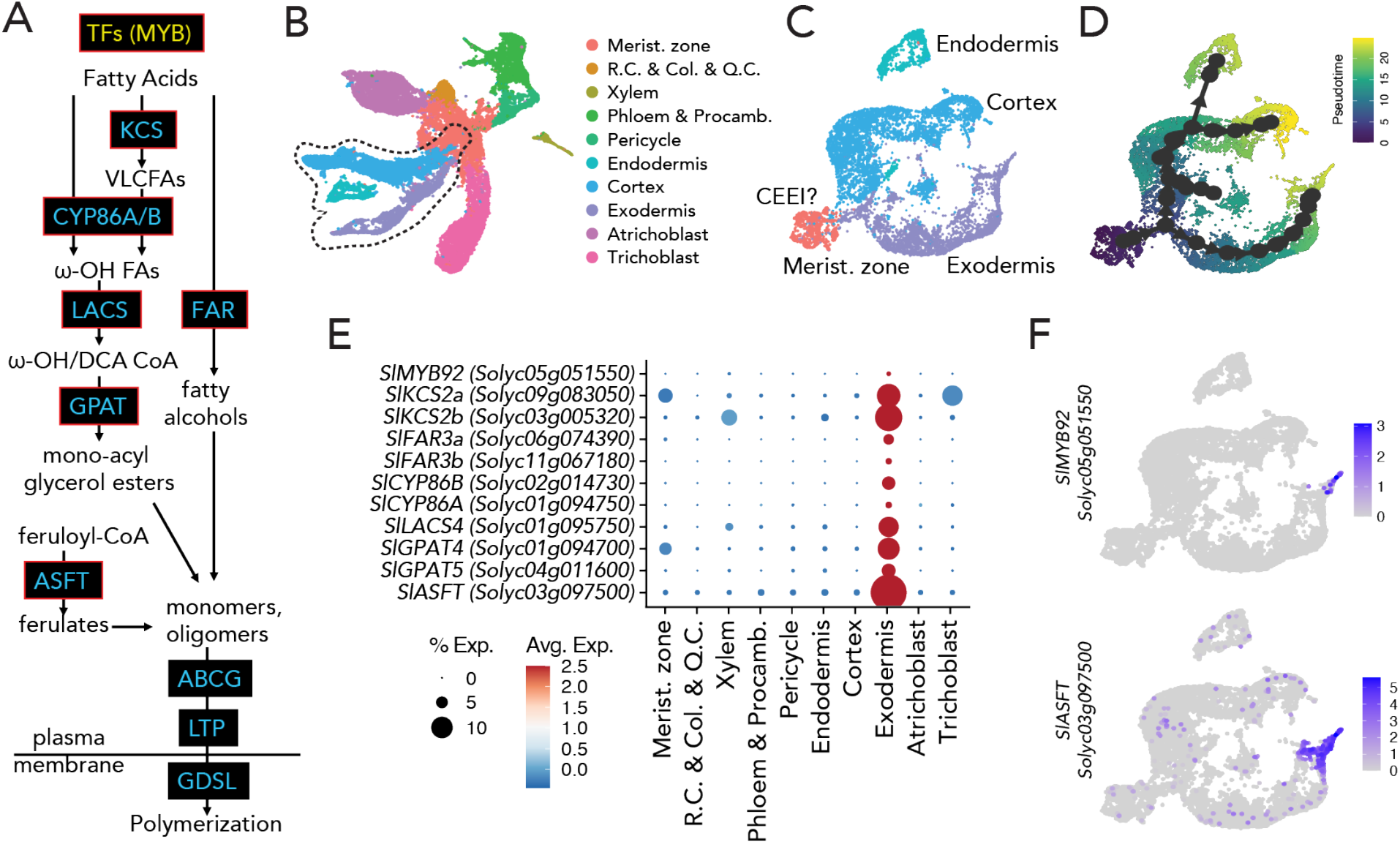
The tomato suberin biosynthetic enzymes and transcriptional regulator are expressed in the mature exodermis. (A) Simplified diagram of the suberin biosynthesis pathway. Boxes indicate gene families involved in each step of the pathway (blue and yellow indicate biosynthetic enzymes and transcriptional regulators, respectively). Genes targeted in this study are outlined in red. (B) Annotated single cell clusters from tomato root 3 cm tip displayed by an integrated uniform manifold approximation and projection (UMAP). (C) UMAP of Cortex/Endodermis/Exodermis-annotated cells that were extracted from the general projection and re-embedded. A small cluster of cells from the meristematic zone clusters were included to help anchor pseudo-time estimations. (D) A pseudo-time trajectory analysis for the cortex/endodermis/exodermis cell populations. (E) Cell type or tissue-specific expression profiles for suberin biosynthetic pathway genes. Dot diameter represents the percentage of cells in which each gene is expressed (% Exp.); and colors indicate the average scaled expression of each gene in each developmental stage group with warmer colors indicating higher expression levels. R.C.: Root cap. Q.C.: Quiescent center. Col: Columella. Procamb: Procambium. CEEI: Cortex-exodermis-endodermis initial. (F) Expression of *SlMYB92* and *SlASFT* in the single cell transcriptome data. The color scale represents log_2_-normalized corrected UMI counts.

### A single cell tomato root atlas to map the exodermal suberin pathway

Although translatome profiles exist for the exodermis (Kajala et al., 2021), these data do not provide resolution of the developmental gradient along which suberin is deposited. To refine the candidate suberin-associated gene set, we conducted single cell transcriptome profiling of the tomato root. Cells were isolated from a three-centimeter segment (two biological replicates) of roots from tomato (M82) seedlings, where suberin deposition is observed. Once the data was pre-processed and filtered for low quality droplets, the remaining high-quality transcriptomes of 22,207 cells were analyzed. After normalization, scaling, and dimensionality reduction via PCA, we visualized cells in a 2-dimensional space using uniform manifold approximation projection (UMAP) (**Figure 2B**) and identified 30 distinct clusters (**Supplemental Figure 5**). Cell clusters were then assigned a cell type identity using the following approaches. We first quantified the overlap with existing cell type-enriched transcript sets from the tomato root (Kajala et al., 2021) and marker genes from each of the clusters. An individual cluster was annotated as a specific cell type given the greatest overlap between the two sets and a significant adjusted p-value (<0.01). Then, pseudotime trajectories were mapped using a minimal spanning tree algorithm (Saelens et al., 2019). The tree was rooted in the root meristematic zone (identified in the previous step), and clusters were grouped into 10 cell types to reflect existing biological knowledge for differentiation of the tomato root (**Figure 2B** and **Supplemental Figure 5**). Lastly, genes with previously validated expression patterns in tomato (Bouzroud et al., 2018; Bucher et al., 1997; Bucher et al., 2002; Gómez-Ariza et al., 2009; Ho-Plágaro et al., 2019; Howarth et al., 2009; Jones et al., 2008; Li et al., 2018; Nieves-Cordones et al., 2010; Ron et al., 2014; von Wirén et al., 2000)(Manzano et al. Biorxiv 2022), transcriptional reporters (Kajala et al., 2021) and predicted cell type markers given their function in Arabidopsis (Shahan et al., 2022), were overlaid on the clusters to refine annotation (**Supplemental Figure 5C**).

Given the successful annotation of these cell types, we focused on the mapped developmental trajectories deriving from a presumed Cortex-Endodermal-Exodermal Initial (CEEI) population (**Figure 2C-D**). Within these three associated trajectories, we localized the cells in which the suberin biosynthetic enzymes and putative regulators were highly expressed (**Supplemental Table 3**). Of these, transcripts of *SlASFT (Solyc03g097500)*, two *FAR (FAR3A: Solyc06g074390; FAR3B: Solyc11g067190)*, two *CYP86* (*SlCYPB86A*: *Solyc01g094750*; *SlCYP86B1*: *Solyc02g014730*), two *KCS2 (SlKCS2a: Solyc09g083050* and *SlKCS2b: Solyc03g005320*), two *GPAT (SlGPAT4: Solyc01g094700; SlGPAT5: Solyc04g011600)*, and one *LACS (SlLACS4: Solyc01g095750*) showed restricted expression at the furthest edge of the exodermal developmental trajectory (**Figure 2E-F** and **Supplemental Figures 6–7**). Of the three transcription factors previously noted (*SlMYB41, SlMYB63* and *SlMYB92*), only *SlMYB92* showed specific and restricted expression in cells at the tip of the exodermal trajectory (**Figure 2F** and **Supplemental Figure 7**). Based on the co-expression and cellular trajectory data, these genes served as likely candidates for an exodermal suberin transcriptional regulator and suberin biosynthetic enzymes.

### Knock-out of candidate genes disrupt exodermal suberin deposition

Functional validation of these enzymes was initially performed by CRISPR-Cas9 gene editing using two or three guide RNAs (**Supplemental Table 4**) and were introduced into tomato via *Rhizobium rhizogenes* (hairy root) transformation (Ron et al., 2014) (**Figure 3A**). Deletion-confirmed mutant alleles of these genes were phenotyped for suberin levels using fluorol yellow staining (**Figure 3B** and **Supplemental Figure 8**). Based on the histological phenotyping, all but the *slcyp86b* mutant showed a decrease in suberin (**Figure 3B**). Further confirmation of decreased suberin levels were obtained by suberin monomer metabolic profiling in the *slgpat5, slgpat4, slasft, sllacs* and *slmyb92* mutants (**Supplemental Figure 8**). These included collective reduction of ferulic acid and sinapic acid aromatic components; fatty acids (C20, C22, C24), ω-hydroxyacids (C18:2, C18:1, C20, C24, C26) and α-ω-diacids (C18:2, C18:1, C18, C20, C22). Given their expression in the terminus of the exodermal developmental trajectory (**Figure 2B**), stable transgenic *slasft* and *slmyb92* deletion mutant alleles were generated by transformation with *A. tumefaciens* using the same guide RNAs. Reduction of suberin levels as well as changes in its accumulation over the root developmental trajectory were observed in two independent mutant alleles of each gene (**Figure 3C-D**, **Supplemental Figure 8**). In the case of *slasft*, the significant delay in suberin deposition and changes in monomer composition in the exodermis differs from its ortholog in Arabidopsis, in which the *atasft* mutant presents no defects in either deposition timing or major monomer composition changes outside of the reduction in ferulate content (Andersen et al. 2021; Molina et al. 2009). We also observed disorganization of the lamellar structure in the *slasft-1* mutant, but not in *slmyb92-1* mutant (**Figure 3E**). While reduction of ferulic acids has been found in mutants of *ASFT* orthologs in potato (Serra et al., 2010), this lamellar disorganization has not been reported before.

**Figure 3:**
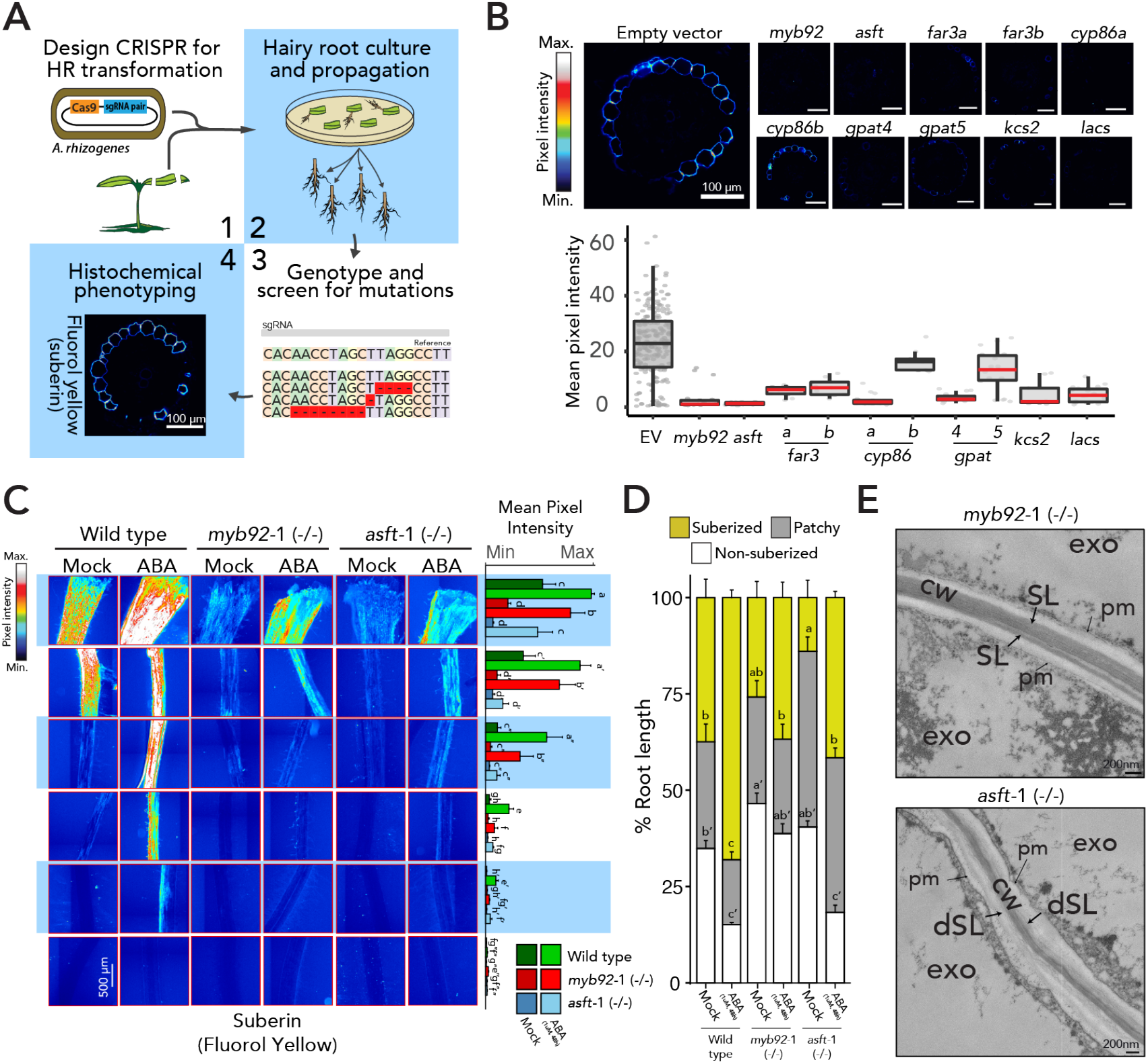
Loss-of-function mutant alleles of candidate genes disrupt exodermal suberization in tomato. (A) Graphical summary of the hairy root (*R. rhizogenes*) mutant screen. (B) Summary of mutant phenotypes of candidate genes in hairy roots. Representative cross sections of mature portions of the roots stained with fluorol yellow (FY) on the top, and overall quantification of fluorol yellow signal across multiple cross sections (n≥6). Red line indicates statistically significant differences in fluorol yellow pixel intensity in the mutant *vs*. wild type as determined with a one-way ANOVA followed by a Tukey HSD test (adj p-value < 0.05). (C) FY staining for suberin in 7-day-old wild-type (repeated from figure 1 for reference), *slmyb92-1* and *slasft-1* plants treated with mock or 1 μM ABA for 48 hours. Whole-mount staining of primary roots across different sections (left) and quantification of fluorol yellow intensity along the root (right) (n ≥ 6). Letters indicate significant differences (one-way ANOVA followed by a Tukey HSD test, adj p-value < 0.05). (D) Developmental stages of suberin deposition in the 7-day-old wild-type and mutant plants treated with mock or 1 μM ABA for 48 h. Zones were classified as non-suberized (white), patchy suberized (gray) and continuously suberized (yellow) (n ≥ 6). (E) Representative transmission electron microscopy cross-sections of *slasft-1* and *slmyb92-1* mutants obtained at 1 mm from the root-hypocotyl junction. The *sl*asft-*1* mutant presents a deficit in suberin lamellar structure. exo = exodermis, SL = suberin lamellae, dSL = defective suberin lamellae, cw = cell wall, pm = plasma membrane.

In Arabidopsis, ABA application increases suberin levels, and four MYB transcription factors are redundantly and partially required for induction of ABA-mediated suberin accumulation in Arabidopsis (AtMYB41, AtMYB53, AtMYB92 and AtMYB93) (Barberon et al., 2016; Shukla et al., 2021). Given the ABA-inducibility of suberin in Arabidopsis and tomato (**Figure 1C-D**) (Barberon et al., 2016; Baxter et al., 2009; Hosmani et al., 2013) we hypothesized that SlMYB92 or SlASFT are necessary for ABA-induced suberin biosynthesis. Therefore, we treated *slmyb92 and slasft* roots with 1μM ABA for 48 hours, a concentration of ABA that is sufficient to increase the completely suberized zone in tomato without perturbing root length (**Figure 1A, Supplemental Figure 10**). Although suberin levels were increased upon ABA treatment in these mutant backgrounds, both in the magnitude of the fluorol yellow signal and the proportion of the root which is completely suberized, the degree to which they were increased is reduced compared to wild type (**Figure 3C-D**). This decrease in suberin levels in the *slmyb92-1* single mutant in both control and ABA-treated conditions is qualitatively greater than what was observed in the single mutant in Arabidopsis (Shukla et al., 2021). The lack of this phenotype in the Arabidopsis single mutant and the in the ABA-induced *atmyb92-1* mutant is explained by redundancy of the A*tMYB41*, AtMYB53, AtMYB92 and AtMYB93 transcription factors (Shukla et al. 2021). We explored whether such redundancy exists in tomato in the hairy root loss-of-function mutant alleles of *slmyb41* and *slmyb63* and whether they were sufficient to decrease exodermal suberin in control and water deficit conditions. ABA treatment was not able to induce suberin to wild type levels in any of the mutants (**Supplementary Figure 8D**). Since the exodermis is first lignified and then suberized (Kajala et al. 2021), we also explored whether SlMYB92, SlMYB41 or SlMYB63 were involved in lignification of the tomato root. However, the levels of lignin in the exodermis and endodermis remained unaffected in any of the hairy root mutants of these transcriptional regulators (**Supplemental Figure 11**), suggesting they do not play a role in the initial lignification of neither the exodermis nor the endodermis.

### Impaired suberin deposition alters the plant’s response to water limitation

Given the suggested link between suberin and drought tolerance, as well as the decreased suberin levels in both control and ABA conditions in our tomato mutants, we hypothesized that the *slmyb92* and *slasft* lines would be more sensitive to water limited conditions compared with wild type plants. We subjected four-week-old well-watered plants to ten days of water deficit conditions (**Methods**). Suberin monomer levels were measured in the root system of *slmyb92-1* and *slasft-1* and WT, in both the water-sufficient and water-limited conditions. Under water-sufficient conditions no significant differences were observed in the very long chain fatty acids, primary alcohols, ω-hydroxy acids, **α**-ω-dicarboxylic acids and aromatic components of suberin (**Figure 4A** and **Supplemental Figure S12**). Under water limitation however, the transcriptional regulator mutant *slmyb92-1* showed a general reduction of most monomer groups compared to wild type. The *slasft-1* mutant, in comparison, was primarily depleted in ferulic acid and its esterification substrates, as well as in individual primary alcohols and ω-hydroxy acids (**Figure 4A**). Furthermore, stem water potential, stomatal conductance, and transpiration rate were significantly decreased in response to water limited conditions in both *slmyb92-1* and *slasft-1* relative to wild type; and leaf relative water content was also decreased in *slmyb92-1* (**Figure 4B-E**). When considering all physiological traits collectively using Principal Component Analysis, *slasft-1* showed a milder water deficit response compared to WT, while *slmyb92-1* was more extreme (**Supplementary Figure 13**). These data demonstrate that decreased suberin levels in the tomato root exodermis directly perturb whole plant performance under water limited, but not water sufficient conditions. Furthermore, changes in specific suberin monomers and the lamellar structure that were observed between the two mutants in response to water limited conditions may differently influence the extent of the physiological response.

**Figure 4:**
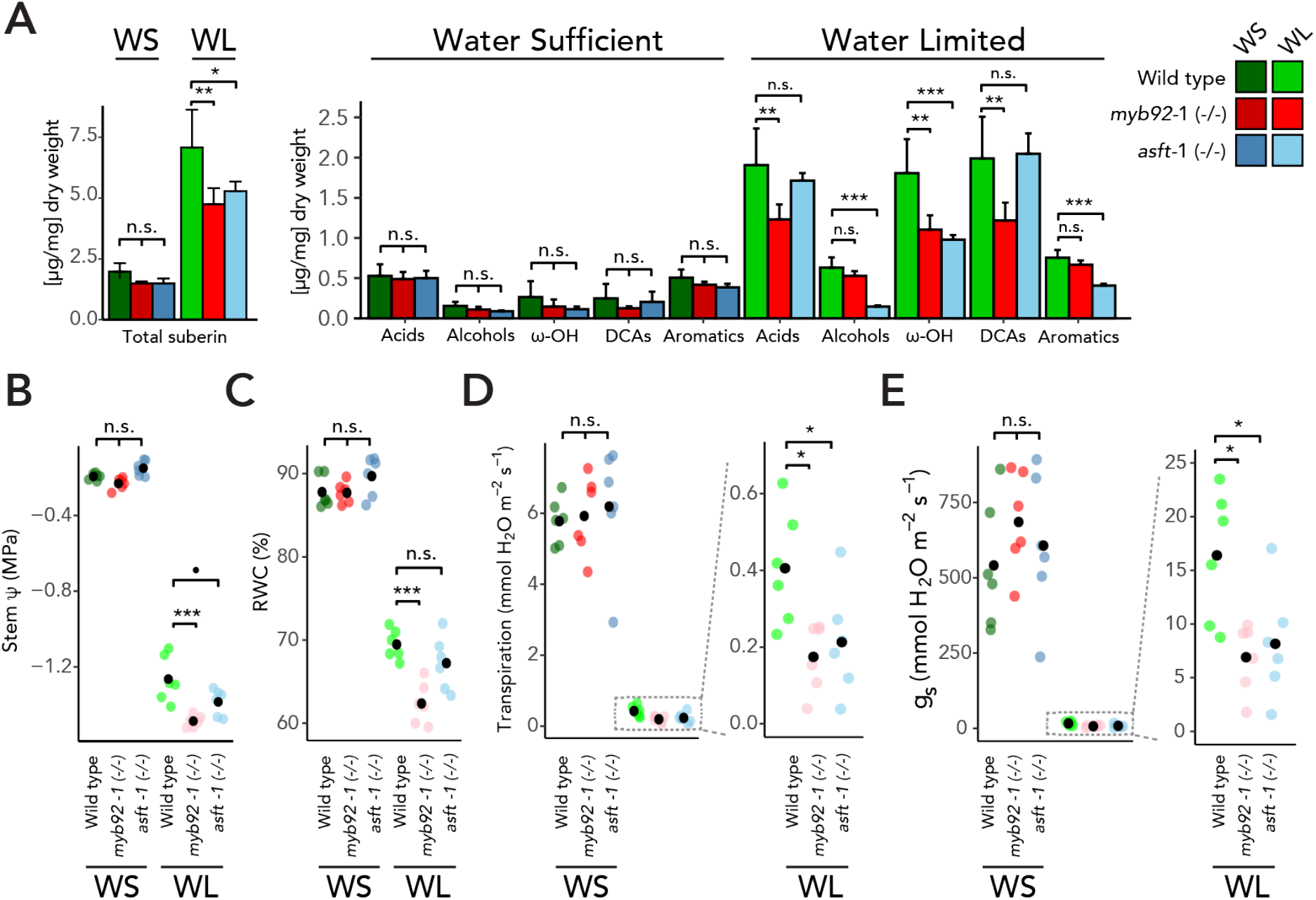
Impaired suberin deposition in *slmyb92-1* and *slasft-1* perturbs their whole plant performances in response to water limitation. (A) Suberin composition in roots of mature one-month-old wild type, *slmyb92-1* and *slasft-1*. Plants were exposed to 10 days of water sufficient (WS) and water limitation (WL) regimes (n = 4, Methods). Acid: fatty acids; Alcohols: primary alcohols; ω-OH: ω-hydroxy fatty acids; DCA: dicarboxylic fatty acid; Aromatics: ferulate and coumarate isomers. Dot plots of recorded values for (B) Stem Water Potential (stem Ψ), (C) Relative Water Content, (D) Transpiration, and (E) Stomatal conductance (g_s_). Dotted line indicates zoom in for better visual resolution of values. Black dots indicate mean values (n=6). One-way ANOVAs for each treatment were performed followed by post-hoc Tukey HSD test. Significance: ‘***’ <0.001 ‘**’ <0.01 ‘*’ <0.05 ‘.’ <0.1.

## DISCUSSION

In the well-characterized Arabidopsis root endodermis, suberin is deposited as a hydrophobic layer between the plasma membrane and the primary cell wall (reviewed in Serra and Geldner, 2022). Developmentally, suberin biosynthesis and deposition occurs as a second step of endodermal differentiation, the first being the synthesis and deposition of the lignified Casparian strip (Naseer et al., 2012). Suberin serves as an apoplastic barrier and a transcellular barrier, thus contributing to the regulation of the movement of water and solutes to the vascular cylinder (Calvo-Polanco et al., 2021). Our collective observations demonstrate that in the tomato root this pathway has distinct spatial regulation relative to Arabidopsis, yet the regulation through developmental time is conserved (**Figure 1**). Spatially, in the tomato root exodermis suberin lamellae are deposited between the exodermal primary cell wall and the plasma membrane all around the cell, similar to the Arabidopsis root endodermis and other suberized apoplastic barriers such as the potato periderm (**Figure 1B**) (Gou et al., 2017; Molina et al., 2009; Serra et al., 2010). In a temporally similar fashion to the Arabidopsis endodermis, there is a non-suberized zone at the root tip, a patchy suberized zone in the middle of the root, and a continuously suberized zone nearer to the root-hypocotyl junction (**Figure 1C-D**).

We obtained clues to the underlying genes controlling exodermal suberin biosynthesis over developmental time by co-expression analysis and single cell transcriptome profiling. Conservation of the biosynthetic pathway between Arabidopsis and tomato is evident from the functional genetic analysis of enzymes within the pathway as well as of the *MYB92* transcription factor. A distinct deviation from this conservation of suberin biosynthesis is the perturbed lamellar structure in the tomato *slasft* mutant. Conversely, in the Arabidopsis and potato mutants, there is a reduction in ferulates associated with suberin, and increases and decreases of a variety of monomers (Molina et al., 2009) (Serra et al., 2010). This suggests both novelty and a conserved function of SlASFT. Although *SlMYB92* was expressed at the end of the exodermal trajectory, the length of this trajectory is likely limited by our ability to completely protoplast cells with secondary cell wall deposition. Indeed, mutants in tomato orthologs of Arabidopsis *MYB41* and *MYB63* showed exodermal suberin phenotypes suggesting these genes may be expressed in later exodermal developmental stages. The decrease of suberin levels in *slmyb92* mutant alleles as measured by fluorol yellow (**Figure 3B-D**) and compositional profiling in hairy roots and stable lines (**Supplemental Figures 8 and 12**) demonstrate likely redundancy in this process. This redundancy in transcriptional regulation, although not necessarily via the same factors, is again conserved between Arabidopsis and tomato (Shukla et al., 2021). What remains to be identified, however, are the factors or regulatory elements that determine exodermal-specific regulation of these enzymes and transcriptional regulators, as well as how they are activated by ABA and why their activity is ABA-independent in *S. pennellii*.

ABA-mediated regulation of suberization is morphologically consistent with what is observed in the Arabidopsis root endodermis, with an increase in both the magnitude of suberin deposition, and an increased proportion of the completely suberized zone (Barberon et al., 2016; Shukla et al., 2021), albeit in a different cell type from the endodermis. At least in the case of the *slmyb92* and *slasft* mutant alleles and the ABA assays, this transcription factor and biosynthetic enzyme influence both developmental and ABA-mediated suberin deposition patterns (**Figure 3C-D**). Further analyses of mutant alleles of the tomato *SlMYB41* and *SlMYB62* transcription factors will determine if a coordinated developmental and stress-inducible regulation of suberin biosynthesis is the norm for exodermal suberin. The degree to which this regulation is dependent on ABA signalling, as it is in Arabidopsis (Barberon et al., 2016), also remains to be observed.

External application of ABA can be considered a proxy for both drought and salt stress response (Zhu 2002; Raghavendra et al. 2010). We tested the necessity of suberized exodermis for whole plant performance under water limited conditions in mature tomato plants (Figure 4). The strongly reduced response of *slmyb92* and *slasft* to ABA was similarly observed upon drought stress. In both experiments *slmyb92-1* and *slasft-1* failed to reach fluorol yellow signals and suberin levels equal to the control. Under control conditions (water-sufficient) we detected overall low suberin levels, which were near the detection limit of 0.003 μg/mg and reduced our ability to identify significant differences between the lines. The effect observed in chemical suberin quantification may have also been attenuated by the sample comprising whole root systems comprised of highly branched lateral roots and including root areas with immature suberin. AtMYB92 is also known to regulate lateral root development in Arabidopsis together with its close ortholog AtMYB93 (Gibbs et al., 2014) and differences in suberin within different root types are a possibility. Regardless, suberin monomeric levels were clearly decreased in the *slmyb92-1* and *slasft-1* mutants in a distinct and overlapping fashion in response to water limited conditions. Consistent with its function, *slasft-1* was primarily defective in accumulation of ferulate, primary alcohols and ω-hydroxyacids (**Figure 4A, Figure S9**); while *slmyb92-1* had defects in fatty acids and the predominant unsaturated C18:1 ω-hydroxyacids and dicarboxylic acids (**Figure 4A, Figure S9**). The more extreme perturbation of physiological responses in response to water limitation in *slmyb92-1* suggests that suberin composed of these fatty acid derivatives play a role in controlling transcellular-mediated uptake of water (**Figure 4B-C**). How the transcellular pathway operates in a root system where this apoplastic barrier is located four cell layers from the vascular cylinder, remains an important and open question.

The role of exodermal suberin as an apoplastic barrier to water flow has been studied in maize and rice, where it was determined as a barrier to water flow, although maize and rice also present a suberized endodermis (Zimmermann et al., 2000). Thus, the role of exodermal suberin alone has never been studied with respect to its influence on plant responses to water limitation. The precise role of endodermal suberin, independent of the Casparian strip, has been studied in Arabidopsis, which lacks an exodermis (Calvo-Polanco et al., 2021). In 21-day-old, hydroponically-grown Arabidopsis plants, the *horst-1, horst-2, horst-1 ralph-1*, pCASP1:*CDEF1* mutants with a functional Casparian strip (Calvo-Polanco et al., 2021), but with reduced suberin (Compagnon et al., 2009; Höfer et al., 2008; Naseer et al., 2012) were monitored for the importance of suberin in water relations. These mutants, except for *horst-2*, have higher Lp_r_ (root hydraulic conductivity) and root aquaporin activity relative to WT (Calvo-Polanco et al., 2021). One can extrapolate that the decrease in stem water potential, transpiration, and stomatal conductance relative to wild type in water limited conditions (**Figure 4B-C**) are a consequence of decreased suberin (*slmyb92*) or perturbations in suberin composition (*slasft*). Assuming our suberin-defective mutants have higher root hydraulic conductivity (Calvo-Polanco et al. (2021)), our hypothesis to reconcile our observations with the higher Lpr would be that our mutants have compromised water use efficiency under water limitation. This could lead to a delayed onset of the drought response such that the water loss is too great to recover by the time stomata are closed. The mechanisms by which this occurs needs to be determined and could benefit from further exploration. The levels of lignin in the exodermis and endodermis were not altered in the mutants of the identified transcriptional regulators (**Supplemental Figure 11**), and perturbations in endodermal lignin alone have no influence on root hydraulic conductivity in Arabidopsis (Calvo-Polanco et al., 2021), thus, lignin plays no role in our observations.

To the best of our knowledge, a plant’s response to water limitation has never been investigated in plants with decreased root exodermal suberin levels. The importance of plant radial and cellular anatomy has also long been known as critical to our understanding of the role of plant roots in water uptake (Steudle, 2000) in the face of water-deficit. Therefore, our findings provide direct evidence, via genetic perturbation, for the role of suberin in a specific cell type mediating tomato’s adaptive response to water-deficit. Further, they impart a model by which exodermal suberin barriers contribute to whole plant water relations, in the absence of a suberized endodermis.

Suberin in plants roots has recently been proposed to be an avenue to combat climate change including via sequestration of atmospheric CO_2_ as well as in conferring drought tolerance (Thompson, 2017). This study provides evidence that root suberin is necessary for tomato’s response to water deficit conditions. Increasing suberin levels within the root exodermis and/or the endodermis may indeed serve as such an avenue. The constitutive production of exodermal suberin in the drought tolerant and wild relative of tomato, *Solanum pennellii* (**Supplemental Figure 3**) (Gur and Zamir, 2004; Gur et al., 2011; Pillay and Beyl, 1990) certainly provides a clue that maintenance of suberin in non-stressed and stressed conditions may result in such a benefit. However, trade-offs of such an increase must also be considered. Increased suberin levels have been associated with pathogen tolerance (Holbein et al., 2019; Kashyap et al., 2022; Thomas et al., 2007), but also can serve as a barrier to interactions with commensal microorganisms (Salas-González et al., 2021), and could constrain nutrient uptake, plant growth or seed dormancy (Beisson et al., 2007; Cohen et al., 2020; To et al., 2020). Regardless, this complex biopolymer serves as an elegant example of how plant evolution has resulted in the different but precise spatiotemporal biosynthesis and deposition patterns of a specialized polymer to enable a plant’s response to the environment.

## MATERIALS & METHODS

### Plant material and growth conditions

All tomato (*Solanum lycopersicum*) lines used in this study were derived from cultivar M82 (LA3475). The *Solanum pennellii* line used was LA0716. Seeds were surface sterilized in 70% (v/v) ethanol for 5 min followed by 50% (v/v) commercial bleach for 20 min and three washes with sterile deionized water. Seeds were plated on 12cmx12cm plates (without sucrose) or in Magenta boxes (with 30 g L−1 sucrose) containing 4.3 g L−1 Murashige and Skoog (MS) medium (Caisson; catalog no. MSP09-50LT), 0.5 g L−1 MES, pH 5.8, and 10 g L−1 agar (Difco; catalog no. 214530) and maintained in a 23°C incubator with 16/8h light/dark periods for 7-10 days, until cotyledons were fully expanded and the true leaves just emerged. At that point, and depending on the experiment, seedlings were either harvested or transferred to soil.

### Tomato transformation

Hairy root transformants were generated based on published work (Ron et al. 2014). In brief, *Rhizobium rhizogenes* (Strain ATCC 15834) transformed with the desired binary vector was used to transform expanding tomato cotyledons. Using a scalpel, 7-10 day old M82 cotyledons were cut and immediately immersed in the bacterial suspension for 20 minutes, blotted on sterile Whatman filter paper, and co-cultivated for 3 days at 25°C in dark on MS agar plates (1X with vitamins, 3% sucrose, 1% agar) without antibiotic selection. Cotyledons were then transferred to MS plates with a broad spectrum antibiotic cefotaxime (200 mg L−1) and kanamycin (100 mg L−1) for selection of successfully transformed roots. Fifteen independent antibiotic-resistant roots were subcloned for each construct for further analysis. Stable transgenic lines were generated by *Agrobacterium tumefaciens* transformation at the UC Davis Plant Transformation Facility.

### Transcriptome Profiling of M82 Roots under Drought and Waterlogging Stress

Seeds of *SlCO2p:TRAP* and *AtPEPp*:TRAP cv. M82 (Kajala et al., 2021) were surface-sterilized with 3% hypochlorite (Clorox) for 20 minutes, rinsed three times with sterile water and plated on 1xMS media with 200 mg/ml kanamycin. Seven days after germination, seedlings were transplanted into 15cm x 15cm x 24cm pots with Turface Athletic Profile Field & Fairway clay substrate (Turface Athletics) pre-wetted with a nutrient water solution (4% nitrogen, 18% phosphoric acid, and 38% soluble potash). Plants were grown in a completely randomized design for 31 days in a Conviron Growth Chamber at 22C, 70% RH, 16/8 hour light/dark cycle and light intensity of 150-200 mmol/m^2^/s. For “well-watered” conditions, we maintained substrate moisture at 40-50% soil water content. We based this selection on pilot experiments where we monitored tomato development and physiology, including photosynthesis measurements with a LICOR 6400XT, local temperature and leaf surface temperatures measured with an infrared digital thermometer, and relative water content (RWC) measured from terminal leaflets from the youngest expanded leaf. As our water deficit treatment, we withheld water from the plants for 10 days prior to harvest, and as our waterlogged condition, we submerged the pot until the root-to-shoot junction. We harvested the roots as close to relative noon as feasible (±2h) by immersing the pot into cool water, massaging the rootball free, rinsing three times sequentially with water, and then dissecting the root tissues and flash-freezing with liquid nitrogen. We harvested the lateral roots at the depth of 6-12 cm and 1-cm root tips of adventitious roots for the RNAseq experiment. RNA sequencing libraries were synthesized from four biological replicates from the *SlCO2p*:TRAP and *ATPEPp*:TRAP lines, with the exception of the control for *SlCO2p*:TRAP lateral roots in control conditions (five biological replicates). Sequencing libraries of adventitious roots were generated for each line in control and waterlogging conditions, and from lateral roots in control, waterlogging and water deficit conditions. Total RNA was isolated from these roots as described in (Reynoso et al., 2019); and non-strand specific random primer-primed RNA-seq library construction performed as described originally by (Townsley et al., 2015). We pooled the RNAseq libraries together and sequenced them with the Illumina HiSeq 4000 at the UC Davis DNA Technologies Core to obtain 50-bp single-end reads.

### RNA-sequencing data processing and analysis - drought, water deficit and introgression line population

RNA-sequencing data processing and analyses for the drought, waterlogging and introgression line population were conducted as previously described (Kajala et al., 2021). Sequences were pooled, and then trimmed and filtered using TrimGalore (Krueger, 2012), with parameter -a GATCGGAAGAGCACA. Trimmed reads were pseudo-aligned to the ITAG3.2 transcriptome (cDNA) (Tomato Genome Consortium, 2012) using Kallisto (v0.43.1) (Bray et al., 2016), with the parameters -b 100–single -l 200 -s 30, to obtain count estimates and transcript per million (TPM) values. Raw RNA-seq read counts were filtered to remove genes with zero counts across all samples. Samples were clustered with cuttreestatic (Langfelder and Horvath, 2008) and outliers removed with a minSize of 10. No outliers were observed for the drought and waterlogging dataset, although GSM2323699 (Toal et al., 2018) from the introgression line dataset was removed as an outlier.

### Generation of tomato CRISPR constructs

Target guides were designed using the CRISPR-PLANT web tool (https://www.genome.arizona.edu/crispr/CRISPRsearch.html) (**Supplemental Table 4**). In cases where CRISPR-PLANT did not specify at least three guides with GC content between 40 to 60%, guides were designed with CRISPR-P V2 (http://crispr.hzau.edu.cn/cgi-bin/CRISPR2/CRISPR), using the U6 snoRNA promoter with < 3 mismatches within the target gene coding sequence. Genomic sequences (ITAG3.2) were retrieved from Phytozome (https://phytozome-next.jgi.doe.gov/) and gene maps were constructed with SnapGene. Primers for genotyping were designed with Primer-BLAST software (https://www.ncbi.nlm.nih.gov/tools/primer-blast/). Primer specificity was checked against *Solanum lycopersicum* RNA entries from NCBI’s Reference Sequence collection, using RefSeq RNA as a database. In cases where only two gRNAs were selected (initial round of the cloning), cloning was performed based on Fauser et al. (2014). In summary, oligo containing the sgRNA PAM sequence were ligated into pMR217/218 vectors, and then recombined via Gateway assembly into a pMR290 vector containing Cas9 and Kan resistance expression cassettes (Bari et al. 2019). In cases where three gRNAs were selected (second round of cloning), cloning of guide RNAs was performed based on Lowder et al. (2015). In summary, oligos containing the sgRNA PAM sequence were phosphorylated and ligated into pYPQ131-3 vectors, and then recombined into pYPQ143 via Golden Gate assembly. A pMR278 vector containing all 3 gRNA expression cassettes was then recombined into a pMR286/289 vector containing Cas9 and Kan resistance expression cassettes (Bari et al. 2019). All the final CRISPR vectors were introduced into *Rhizobium rhizogenes* (Hairy roots) and *Agrobaterium tumefaciens* (Stable lines) for transgenic generation.

### CRISPR-Cas9 mutant generation and analysis

Transgenic hairy root and stable lines containing the CRISPR binary vectors were screened for mutations in the genes of interest. Independently transformed lines were genotyped at the targeted genomic region (small guide RNA and oligo sequences are found in **Supplemental Tables 4 and 5**). In the case of hairy roots, at least 2 lines containing large deletions in both alleles in the gene of interest were kept for further analysis. In the case of stable transformants, first-generation (T0) transgenics were genotyped via Sanger sequencing. After genotyping and self-pollination in a greenhouse, T1 seeds from T0 plants with the mutated genes were sown. Plants were screened, and lines that did not contain the CRISPR construct anymore and had homozygous mutant alleles were selected. T2 and T3 seeds obtained from self-pollination in a greenhouse were used in subsequent experiments.

### Water Deficit Assay

7-day old seedlings were transferred to 0.5L cones containing Turface Athletic Profile Field & Fairway clay substrate (Turface Athletics) that was pre-wetted with a nutrient water solution (containing 4% nitrogen, 18% phosphoric acid, and 38% soluble potash). All pots were weight adjusted, and a small set of pots were dried so that the percentage of water in the soil could be calculated based on pot weight. Plants were then grown in a completely randomized design for three weeks in a Conviron Growth Chamber at 22°C, 70% RH, 16/8 hour light/dark cycle and light intensity around 150 μmol/m2/s, and watered to soil saturation every other day. At the end of the first week in the chamber, vermiculite was added to the top of the cones to limit water evaporation from the soil. Following three weeks, plants of each line were randomly split into two groups and plants were exposed to different treatments for 10 days (drought and control). Six “control” plants per line were kept in a “water sufficient” regime and watered to soil saturation with nutrient solution every day. Six “drought” plants per line were kept in a “water limited” regime as follows: Plants were gradually subjected to water deficit by adjusting pots weight daily with nutrient solution (to the highest weight of the set) until a target soil water content of 40-50% was obtained. On the day of harvesting, between 9:00 to 12:00 am, stomatal conductance and transpiration were measured on the abaxial surface of the terminal leaflet of the 3rd leaf or the youngest fully expanded leaf using LICOR-6400XT. Light intensity was kept at 1,000 μmol m^−2^ s^−1^, with a constant air flow rate of 400 μmol s^−1^ and a reference CO_2_ concentration of 400 μmol CO_2_ mol^−1^ air. The 3rd (either left or right) primary leaflet was collected for measuring relative water content using a modified version of a previously established protocol (Sade et al. 2015). In short, fresh leaves were cut with a scalpel leaving a 1-cm-long petiole and the total fresh weight (TFW) was measured. Leaves were then placed in individual Zipper-locked plastic bags containing 1 mL of deionized water, making sure that only the leaf petiole is immersed in the solution. Bags were incubated at 4 °C. After 8 h, leaves were taken out of the bags and put it between two paper towels to absorb excess water; and then weighed to determine the turgid weight (TW). Each sample was then placed into a paper bag and dried in a 60 °C dry oven for 3-4 days. Dried samples were weighed to determine the dry weight (DW), and relative water content was calculated as: RWC(%)= (TFW-DW)*100/(TW-DW). A section of the 4th leaf, containing the terminal and primary leaflets was used to measure stem water potential (McCutchan and Shackel, 1992) using a pump-up pressure chamber (PMS Instrument Company, Albany, OR). The root systems were harvested by immersing the cone into water, massaging the root ball free, and rinsing with water to remove as much clay substrate as possible. Plants were then placed on paper towels to remove excess water, and the middle section of the root system was sectioned using a scalpel. Around 300mg of the dissected root tissue were added to Ankon filter bags (sealed with a staple). Bags were transferred into a glass beaker, an excess of chloroform:methanol (2:1, v/v) was added and extracted for 2h. Fresh chloroform:methanol (2:1, v/v) was replaced and the extraction was repeated overnight under gentle agitation (twice). Fresh chloroform:methanol (1:2, v/v) was added and extracted for 2h. The extraction was repeated overnight twice with fresh chloroform:methanol (1:2, v/v). Finally, samples were extracted with methanol for 2h. Methanol was removed, and bags were dried in a vacuum desiccator for 72h. Suberin monomer analysis was performed in these samples as stated below.

### ABA Assay

Seedlings were germinated in MS plates as stated above. 5 days after germination, seedlings from a plate were randomly transferred to fresh MS plates containing either 1 μM ABA or mock. After 48h of treatments in a 23°C incubator with 16/8h light/dark, roots were harvested and used in subsequent analyses.

### Co-expression network analysis

Co-expression network modules were generated with the WGCNA R package (v1.70). Bulk RNAseq libraries from M82 roots under drought stress and the control, and a published root expression dataset from an IL panel (Toal et al. 2018) were used for this analysis. Libraries were quantile normalized and a soft threshold of 8 was used to create a scale-free network. A signed network was created choosing a soft thresholding power of 8, minModuleSize of 30, the module detection sensitivity deepSplit of 2, mergeCutHeight of 0.3. Genes with a consensus eigengene connectivity to their module eigengene lower than 0.2 were removed from the module (minKMEtoStay). Modules were correlated with upregulated genes in DCRi lines from the Lashbrooke et al. 2016 paper using Fisher’s exact test.

### Protoplast isolation and scRNA-seq

This protocol is a modified version of the Arabidopsis single cell sequencing in Shahan et al. (2021). In summary, seven days after sowing, 50-100 primary roots per sample of length ~3 cm from the root tip were cut and placed in a 35 mm-diameter dish containing a 70 μm cell strainer and 4.5 mL enzyme solution (1.25% [w/v] Cellulase R10, 1.25% Cellulase RS, 0.3% Macerozyme R10, 0.12% Pectolyase, 0.6 M mannitol, 20 mM MES (pH 5.7), 20 mM KCl, 10 mM CaCl2, 0.1% bovine serum albumin, and 0.000194% (v/v) - mercaptoethanol). Cellulase Onozuka R10, Cellulase Onozuka RS, and Macerozyme R10 were obtained from Yakoult Pharmaceutical Industries. Pectolyase was obtained from Sigma-Aldrich (P3026). After digestion at 25°C for 2 hours at 85 rpm on an orbital shaker with occasional stirring, the cell solution was filtered twice through 40 μm cell strainers and centrifuged for 5 minutes at 500 x g in a swinging bucket centrifuge with the acceleration set to minimal. Subsequently, the pellet was resuspended with 1 mL washing solution (0.6 M mannitol, 20 mM MES (pH 5.7), 20 mM KCl, 10 mM CaCl2, 0.1% bovine serum albumin, and 0.000194% (v/v) - mercaptoethanol) and centrifuged for 3 minutes at 500 x g. The pellet was resuspended with 1mL of washing solution and transferred to a 1.7mL microcentrifuge tube. Samples were centrifuged for 3 minutes at 500 x g and resuspended to a final concentration of ~1000 cells/μL. Single cell transcriptome libraries were then prepared by the UC Davis DNA Technologies Core. The protoplast suspension was then loaded onto microfluidic chips (10X Genomics) with v3 chemistry to capture 10,000 cells/sample. Cells were barcoded with a Chromium Controller (10X Genomics). mRNA was reverse transcribed and Illumina libraries were constructed for sequencing with reagents from a 3’ Gene Expression v3 kit (10X Genomics) according to the manufacturer’s instructions. Sequencing was performed with a NovaSeq 6000 instrument (Illumina) to produce 100bp paired end reads.

### Protoplasting-Induced Genes

Protoplast samples were obtained following the same strategy as for the single cell library preparation. Once protoplasts were purified, total RNA was extracted using the Direct-zol RNA Miniprep kit (ZYMO). Bulk RNAseq libraries were prepared using the QuantSeq 3’ mRNA-Seq Library Prep Kit FWD (Lexogen). Barcoded libraries were then pooled together, and PE 150-bp reads were sequenced on the NovaSeq 6000 instrument (Illumina) at the UC Davis DNA Technologies Core. Sequences were pooled, and then trimmed and filtered using Trim Galore! (v0.6.6). R1 Trimmed reads were pseudo-aligned to ITAG4.1 transcriptome (cDNA) using Kallisto (v0.46.2), with the parameter -b 100, to obtain count estimates and transcript per million (TPM) values. Differential expression analysis was performed in edgeR (v3.34.1). Differentially expressed genes with adj.P.Val < 0.05 and logFC > 2 were selected as protoplast-induced genes.

### Single Cell Transcriptome Analysis

FASTQ files were mapped using cellranger (10X Genomics). Reads were aligned to the tomato genome (SL4.0) with the ITAG4.1 gene annotation file. Organellar (mitochondria and plastid) sequences and gene annotation were appended to the main genome and annotation files. Protoplasting-induced (**Supplemental Table 2**) genes were removed. Genes with counts in 3 cells or less were also removed. Low quality cells that contained less than 500 unique molecular identifiers (UMIs) were filtered out. Additionally, cells with >1% UMI counts belonging to organelle genes were filtered out. Data was then normalized using Seurat (v4.0.5) (Satija et al., 2015), followed by principal component analysis (PCA) and non-linear dimensionality reduction using UMAP. Fifty principal components were calculated and UMAP embedding was generated using the initial 35 principal components. Cluster-enriched genes were computed using the “FindAllMarkers” function in Seurat using the only.pos = TRUE, min.pct = 0.1, logfc.threshold = 0.25 parameters.

### Single cell cluster annotation

The clusters were annotated based on the overlap of cluster marker genes and a set of cell type-enriched marker genes from (Kajala et al., 2021). Given a set of tissue specific markers for T number of tissue types we call these sets M_i_ (i=1…A), with Mi = {m_i1_, m_i2_… m_in_}, m_ij_ representing genes in the cell type-enriched marker list. These tissue specific markers are mutually exclusive such that no genes appear in two different sets (M_ij_ ^1^ M_km_ for any i, k). We first identified the marker genes from Seurat-generated cluster markers Si = {s_i1_, s_i2_, … s_in_}, (i=1…C), where C equals the number of Seurat generated clusters. We generated an overlapping table between Mi markers and Si markers which we represent in the table as Tij (i=1…T, j=1…C). For each Seurat cluster, we hypothesize that the cells with the highest number of overlapping markers Tij is the cell type of this cluster. A chi-squared test was used to determine the statistical significance of the maker overlap using the following formula:

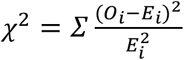

With i=1,2 and

O1 = number of highest overlapping markers argmax(i)T_ij_

E1 = expected number of overlapping markers sum (T_ij_)/N, N = number of tissue types.

O2 = sum of markers that overlap with all other clusters sum (T_ij_) i^1^imax

E2 = expected number of markers that overlap with all other clusters

A Bonferroni-corrected p value was used to select significant marker overlaps. This process was repeated for the second highest and third highest overlapping markers until the corrected p value is higher than 0.01. An individual cluster was assigned the annotation of the tissue types that had the most genes overlapping between the two marker sets, provided the adjusted p value was significant for the overlap.

### Trajectory Analysis of Exodermis Cell types

A trajectory analysis was run for the ground tissue cells (likely exodermis, cortex and endodermal cells derived from the cortex-endodermis initial (Ron et al., 2013) cells after selecting and re-clustering the cell types annotated as exodermis and meristematic zone (clusters 0, 3, 8, 11, 12, 14, 15, 23, 25, 28). Trajectory analysis was performed using dynverse (Saelens et al., 2019) and tidyverse in R. Gene expression matrices, dimensionality reduction and clustering were imported into the dynverse wrapper from Seurat and a starting cell was decided within the meristematic zone cluster and trajectory inference was run using the minimum spanning tree (MST) algorithm. The MST method and UMAP co-ordinates from Seurat were used as input for mclust (Scrucca et al., 2016) in R. Predictive genes or genes that were differentially expressed along the trajectory, specific branches and milestones were identified and visualized with a heatmap using dynfeature within the R package dynverse.

### Histochemical Analysis

Root suberin was observed after Fluorol Yellow (FY) staining as described in (Lux et al., 2005). For sections, roots were divided in 1 cm segments, embedded in 4% agarose, and sliced in 120 um sections using a vibratome. Sections were then incubated in FY088 (0.01%w/v, dissolved in lactic acid) for 1 hour at RT in darkness, rinsed three times with water, and counterstained with aniline blue (0.5% w/v, dissolved in water) for 1 hour at RT in darkness. Confocal Laser Scanning microscopy was performed on a Zeiss Observer Z1 confocal with the 20X objective and GFP filter (488nm excitation, 500-550nm emission). For whole roots, suberin was observed in seven-day-old *S. lycopersicum* wild type or mutant seedlings. Whole roots were incubated in methanol for 3 days, changing the methanol daily. Once cleared, roots were incubated in Fluorol Yellow 088 (0.05%w/v, dissolved in methanol) for 1 hour at room temperature in the dark, rinsed three times with methanol, and counterstained with aniline blue (0.5% w/v, dissolved in methanol) for 1 hour at room temperature in the dark. Roots were mounted and observed with the EVOS cell imaging system (ThermoFisher) using the GFP filter (488nm excitation, 500-550nm emission). Root sections were also stained with basic fuchsin (Fisher scientific Cat no 632-99-5). 1 cm segments from the root tip were embedded in 3% agarose and were sectioned at 150-200 μM using a vibratome (Leica VT1000 S). The sections were stained in Clearsee buffer (Ursache et al. 2018) with basic fuchsin for 30 minutes and then washed two times. Confocal Laser Scanning microscopy was performed on a Zeiss LSM700 confocal with the 20X objective, basic fuchsin: 550-561 nm excitation and 570-650 nm detection.

### Suberin Monomer Analysis

Four biological replicates were analyzed for each genotype. An average of 80 mg fresh weight root tissue per biological replicate was washed and immediately placed in a 2:1 solution of chloroform:methanol. Subsequently, root samples were extracted in a Soxhlet extractor for 8 h, first with CHCl_3_, afterwards with methanol to remove all soluble lipids. The delipidated tissues were dried in a desiccator over silica gel and weighed. Suberin monomers were released using boron trifluoride in methanol at 70°C overnight. Dotriacontane was added to each sample at a concentration of 0.2 μg μl^−1^ as an internal standard, saturated NaHCO_3_ was used to stop the transesterification reaction, and monomers were extracted with CHCl_3_. The CHCl_3_ fraction was washed with water, and residual water removed using Na_2_SO_4_. The CHCl_3_ fraction was then concentrated down to ~50 μl, and derivatized with N,N-bis-trimethylsilyltrifluoroacetamide (BSTFA) and pyridine at 70°C for 40 minutes. Compounds were separated using gas chromatography (GC) and detected using a flame ionization detector (FID; 6890N Network GC System; Agilent Technologies, Santa Clara, CA, USA) basically as described in (Franke et al., 2005). Compound identification was accomplished using an identical gas chromatography system paired with a mass spectroscopy selective detector (GC-MS; 5977A MSD; Agilent Technologies). Compounds were identified by their characteristic fragmentation spectra pattern with reference to an internal library of common suberin monomers and the NIST database.

### Transmission Electron Microscopy

Tomato roots were fixed in 2.5% glutaraldehyde solution (EMS, Hatfield, PA) in phosphate buffer (PB 0.1 M [pH 7.4]) for 1 hour at room temperature and subsequently fixed in a fresh mixture of osmium tetroxide 1% (EMS) with 1.5% potassium ferrocyanide (Sigma, St. Louis, MO) in PB buffer for 1 hour at room temperature. The samples were then washed twice in distilled water and dehydrated in acetone solution (Sigma, St Louis, MO, US) in a concentration gradient (30% for 40 minutes; 50% for 40 minutes; 70% for 40 min and 100% for 1 hour 3 times. This was followed by infiltration in LR White resin (EMS, Hatfield, PA, US) in a concentration gradient (33% LR White 33% in acetone for 6 hours; 66% LR White in acetone for 6 hours; 100% LR White for 12 hours two times) and finally polymerized for 48 hours at 60°C in an oven in atmospheric nitrogen. Ultrathin sections (50 nm) were cut transversely at 2, 5 and 8 mm from the root tip, the middle of the root and 1 mm below the hypocotyl-root junction, using a Leica Ultracut UC7 (Leica Mikrosysteme GmbH, Vienna, Austria), picked up on a copper slot grid 2×1mm (EMS, Hatfield, PA, US) and coated with a polystyrene film (Sigma, St Louis, MO, US). Micrographs and panoramic images were taken with a transmission electron microscope FEI CM100 (FEI, Eindhoven, The Netherlands) at an acceleration voltage of 80kV with a TVIPS TemCamF416 digital camera (TVIPS GmbH, Gauting, Germany) using the software EM-MENU 4.0 (TVIPS GmbH, Gauting, Germany). Panoramic images were aligned with the software IMOD (Kremer et al, 1996)(Kremer et al., 1996).

### Phylogenetic tree construction

Phylogenetic trees were generated using the methods described in (Kajala et al., 2021). Blastp (Madden, 2003) was used to identify homologous sequences from forty-two proteomes with options “-max_target_seqs 15 -evalue 10E-6 - qcov_hsp_perc 0.5 -outfmt 6”. A multiple sequence alignment was generated with MAFFT v7 (option-auto) (Katoh and Standley, 2013), trimmed with trimal (Capella-Gutierrez et al., 2009) with the setting “-gappyout” and a draft tree generated with FastTree (Price et al., 2010). Tree construction methodology was informed by (Rokas, 2011). To generate a maximal likelihood phylogenetic tree, RAxML was used with the option -m PROTGAMMAAUTO and bootstrapped 100 times. Edges with less than 25% bootstrapped support were collapsed using TreeCollapserCL4 (http://emmahodcroft.com/TreeCollapseCL.html).

## SUPPLEMENTAL FIGURE LEGENDS

**Figure S1.**
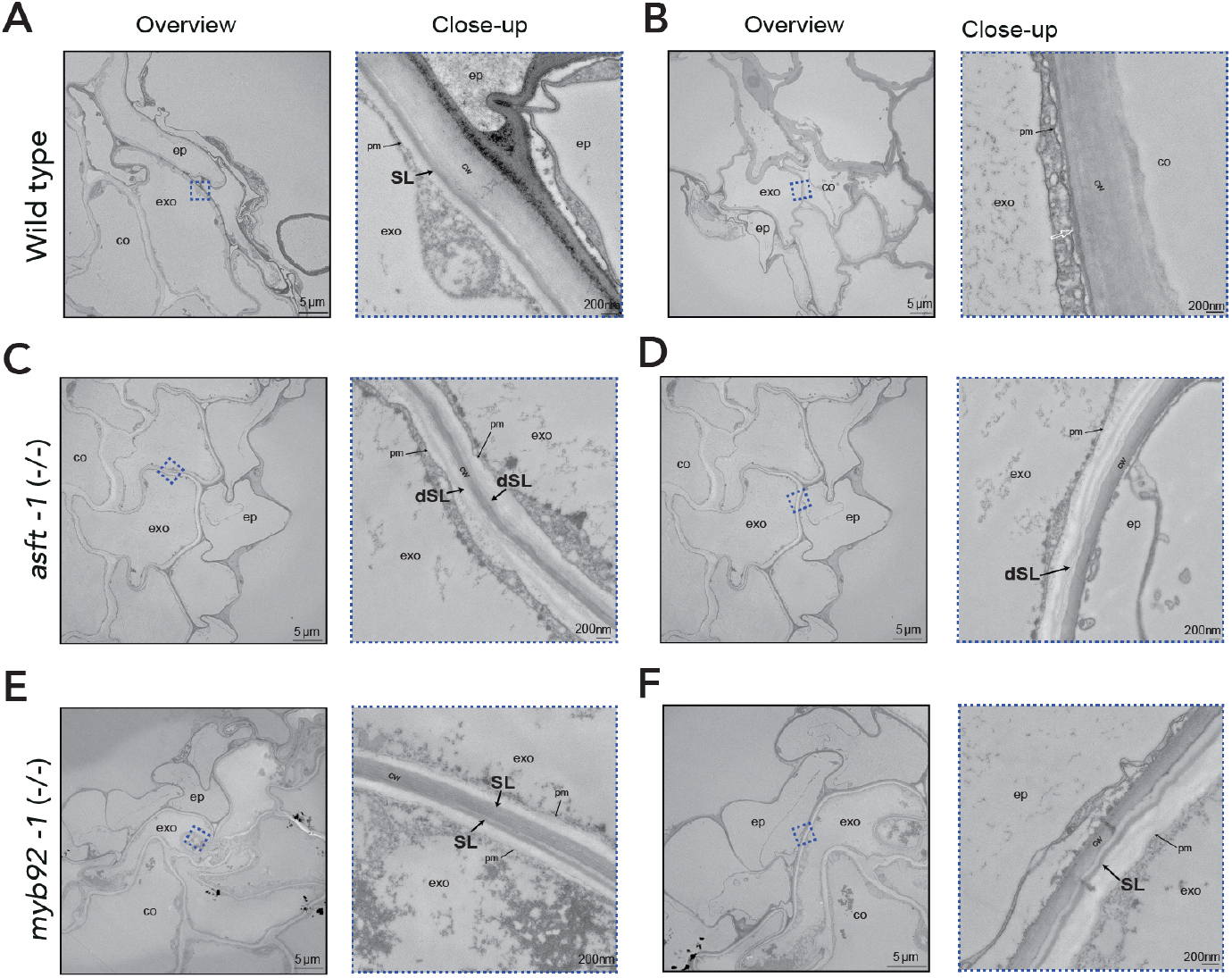
Suberin deposition and ultra-structure in tomato exodermal cells. Transmission electron microscopy cross-sections of 7-day-old wild type, *asft-1* and *myb92-1* plants obtained at 1 mm from the root-hypocotyl junction. Overview shows the epidermal, exodermal and inner cortex layers. Close-up (zone defined with blue dashed lines) shows the presence or absence of suberin lamellae. Black arrows indicate the presence of suberin lamellae, white arrow indicates areas where suberin lamellae could not be detected. scale bars = 5 μm for overview and 200nm for close-up. Close-ups of C & E are repeated in Figure 3. co = cortex, exo = exodermis, ep = epidermis, SL = suberin lamellae, dSL = defective suberin lamellae, cw = cell wall, pm = plasma membrane.

**Figure S2.**
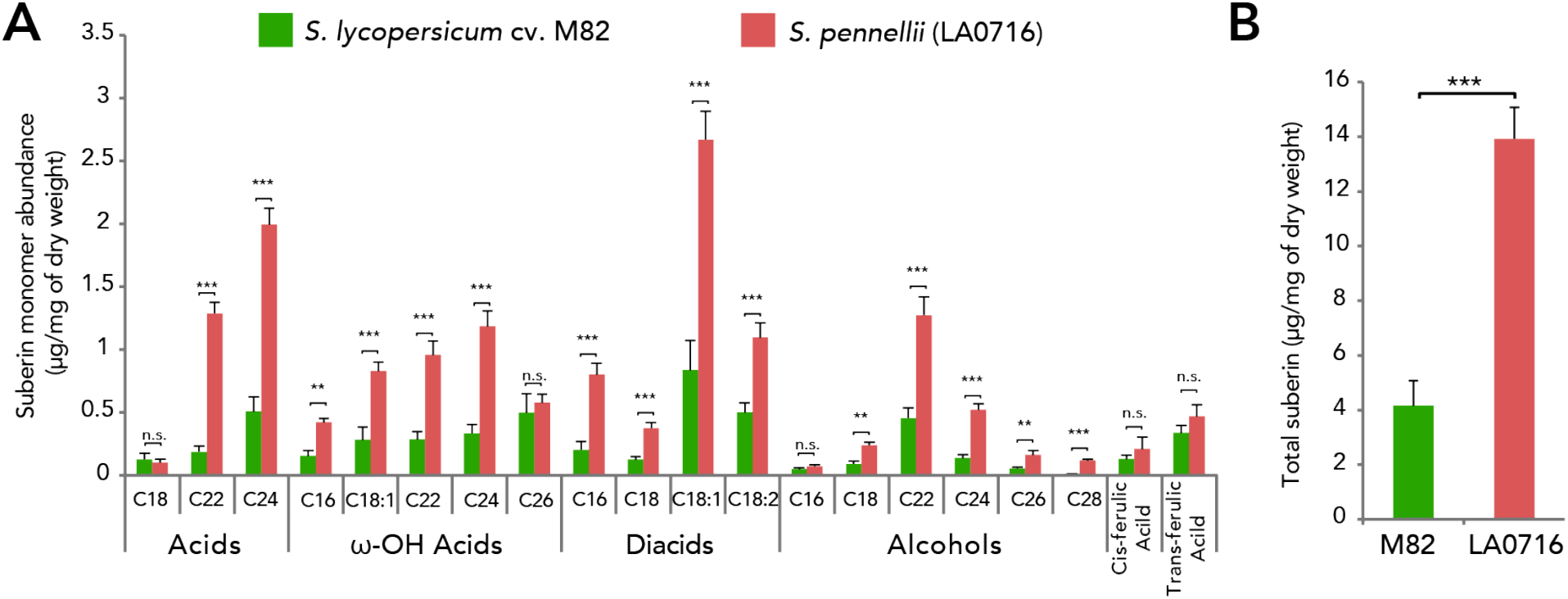
Roots of tomato wild relative *Solanum pennellii* (LA0716) contain significantly more suberin than *Solanum lycopersicum* (cv. M82). (A) Breakdown by monomer abundance. Plants were grown in MS plates and collected 7 days after sowing (n = 5, Methods). Acid: fatty acids; Alcohols: primary alcohols; ω-OH: ω-hydroxy fatty acids; DCA: dicarboxylic fatty acid; Aromatics: ferulate and coumarate derivatives. Signif. codes: ‘***’ <0.001 ‘**’ <0.01, n.s.: “not significant”. (B) Total abundance of suberin expressed as μg mg^−1^ of total dry weight.

**Figure S3.**
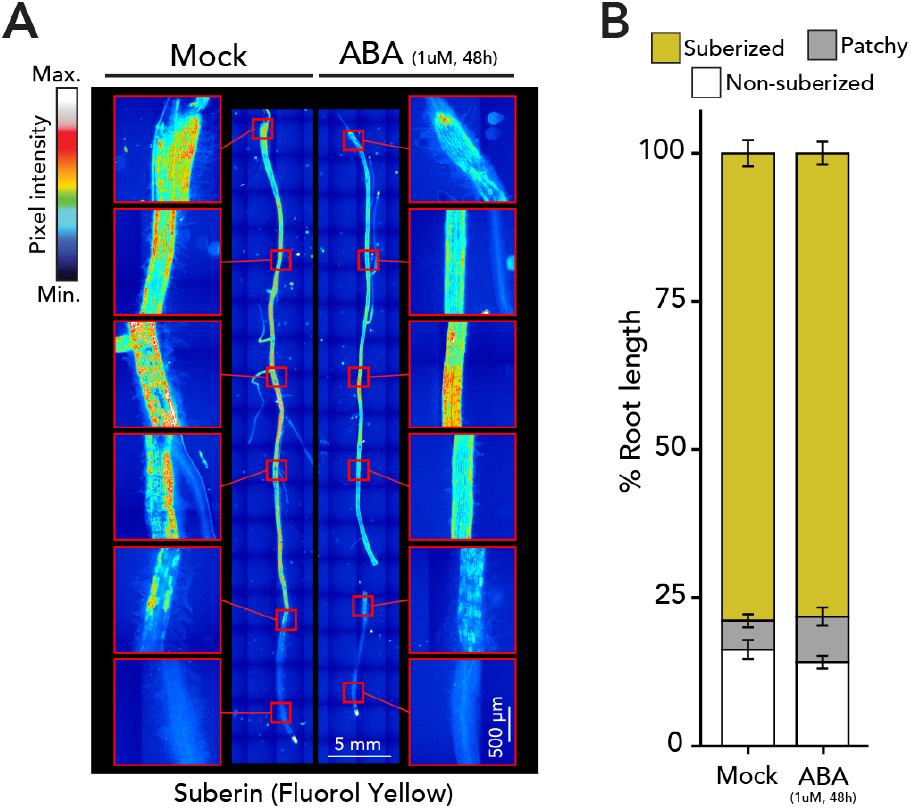
Suberin levels remain unchanged in response to 1 μM ABA in *Solanum pennellii*. (A) Fluorol yellow staining for suberin in tomato wild relative *S. pennellii* (LA0716) 7-day-old plants treated with mock or 1 μM ABA for 48 h. Whole-mount staining of primary roots. (B) Developmental stages of suberin deposition of plants treated with mock or 1 μM ABA for 48 h. Zones were classified in non-suberized (white), patchy suberized (gray) and continuously suberized (yellow), n = 7, error bars: SD.

**Figure S4.**
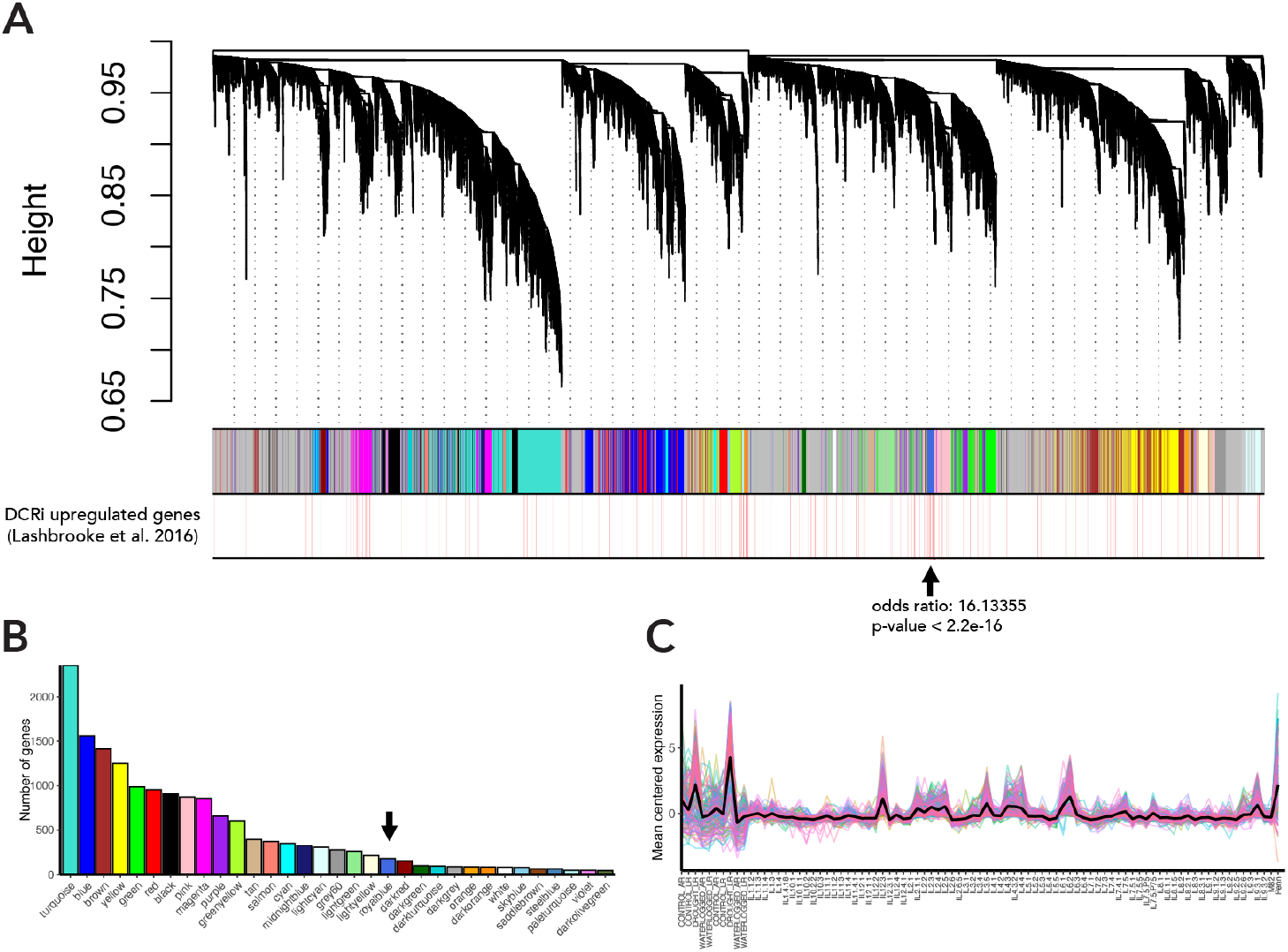
WGCNA analysis of drought and IL datasets identifies a module enriched in suberin-related genes. (A) Gene dendrogram obtained by average linkage hierarchical clustering. The different colors underneath the dendrogram show module assignment as determined by the Dynamic Tree Cut. The bottom panel highlights (marked as thin red lines) the genes identified as upregulated in the suberin overexpression line DCRi in Lashbrooke et al. 2016. The “royal blue” (black arrow) module was significantly enriched for DCRi-upregulated genes. (B) Sizes of all gene modules identified in the analysis. The “royal blue” module (black arrow) contained 180 genes. (C) Mean centred gene expression of all the members of the “royal blue” module. Peaks in expression can be found in drought samples, in *Solanum pennellii* and in certain introgression lines.

**Figure S5.**
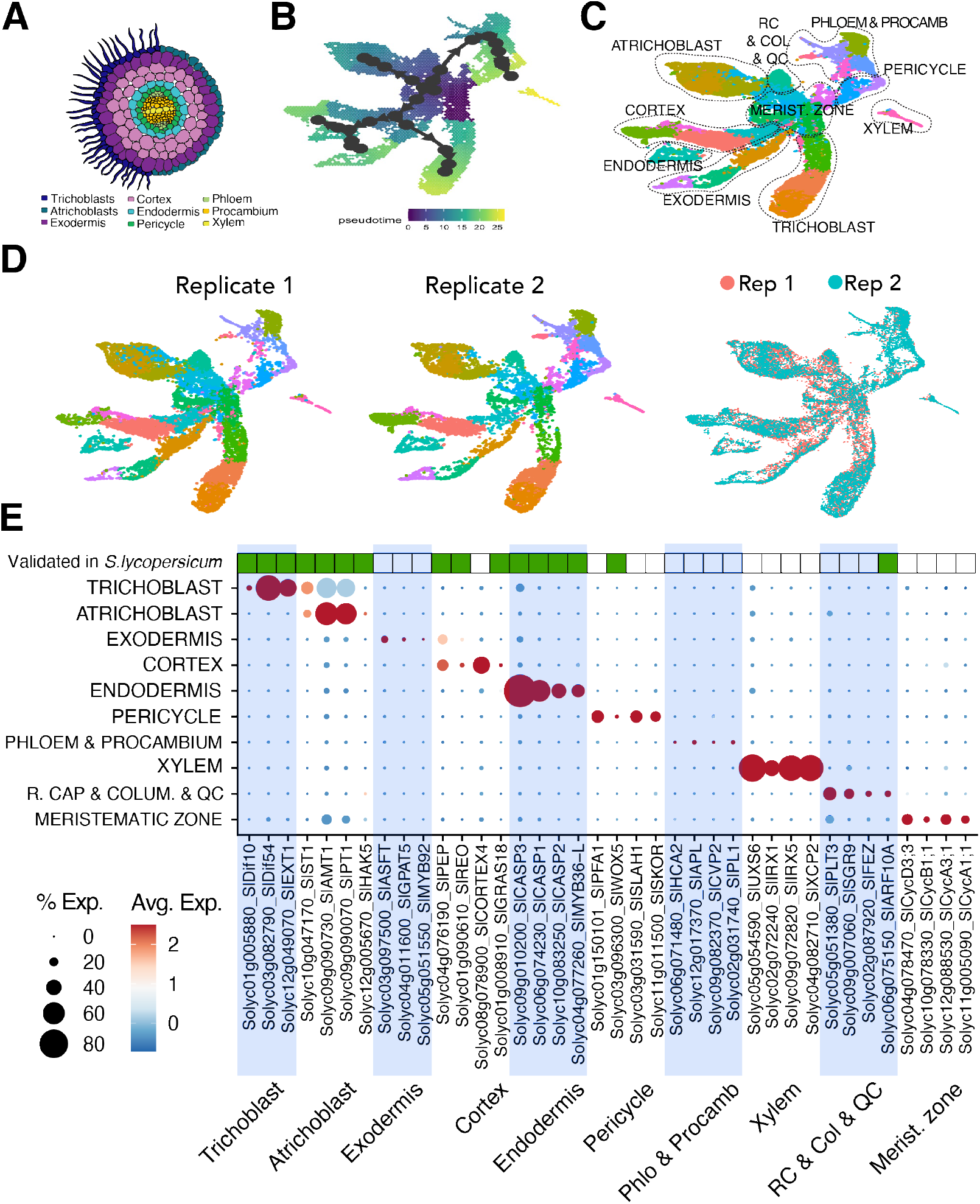
Single cell transcriptome atlas of the tomato root. (A) Graphical depiction of a tomato root section with cell types profiled in the single cell population. (B) A pseudo-time trajectory analysis ran on the population. (C) Annotation of single cell clusters displayed by an integrated uniform manifold approximation and projection (UMAP). Circles indicate subpopulations clustered together. (D) Reproducibility of biological replicates as observed by UMAP and cluster identification. (E) Expression profiles for 39 genes expressed across the major root tissue types. Dot diameter represents the percentage of cells in which each gene is expressed (% Exp.); and colors indicate the average scaled expression of each gene in each developmental stage group with warmer colors indicating higher expression levels. Top row indicates whether the gene’s expression has been validated in *S. lycopersicum* previously published work. R.C.: Root cap. Q.C.: Quiescent center. Col: Columella. Procamb: Procambium. Phlo: Phloem.

**Figure S6.**
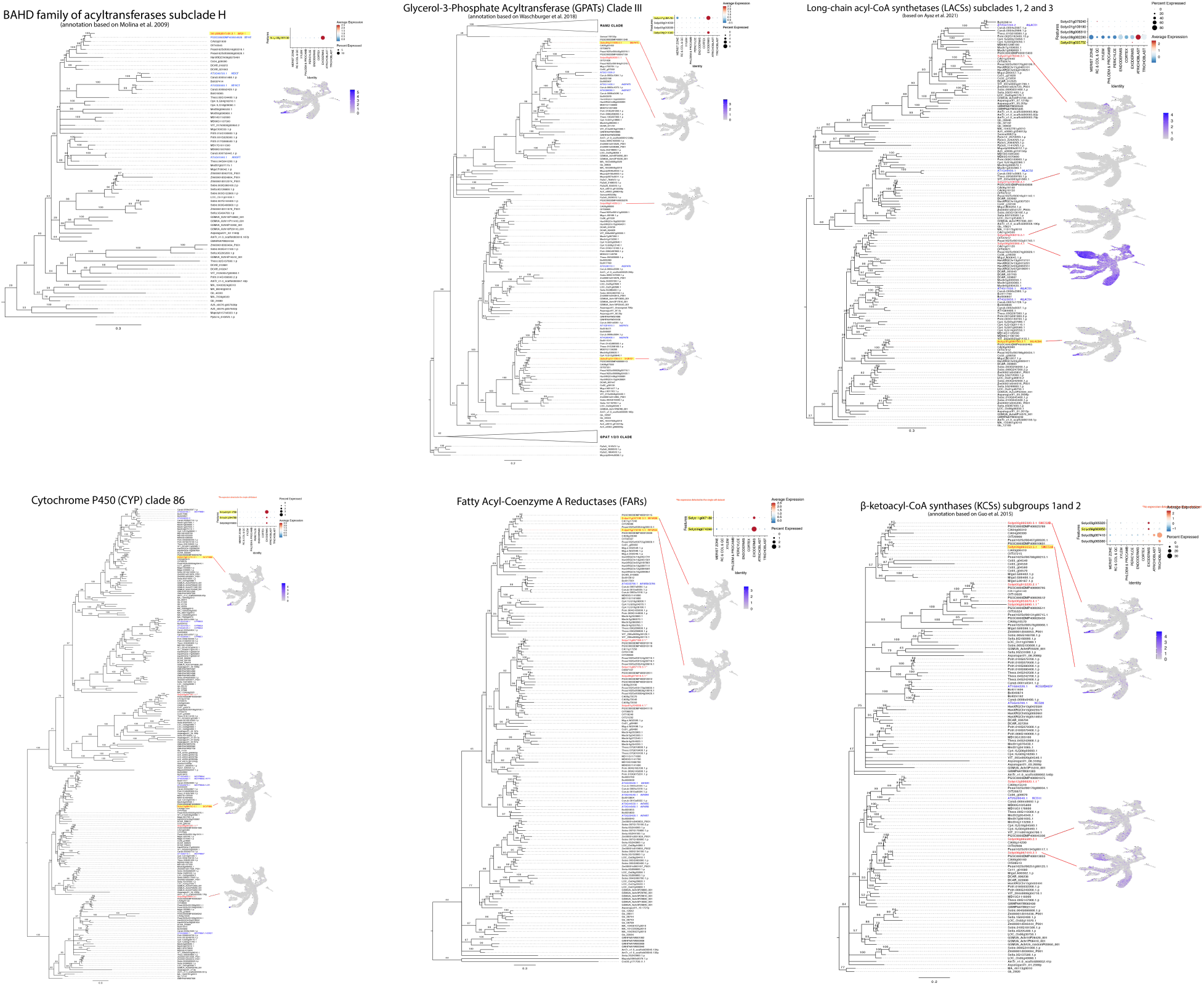
Phylogenetic tree and single cell expression of potential suberin biosynthesis genes. (Left) Phylogenetic trees generated using protein sequences of several plant species. *S. lycopersicum* genes are highlighted in red. *A. thaliana* genes are highlighted in blue for reference. AmTr: *Amborella trichopoda*, AT: *Arabidopsis thaliana*, Asparagus: *Asparagus officinalis*, Azfi: *Azolla filiculoides*, Bol: *Brassica oleracea*, Carub: *Capsella rubella*, CA: *Capsicum annuum*, Cc: *Coffea canephora*, Cp: *Cucurbita pepo*, DCAR: *Daucus carota*, Gb: *Ginkgo biloba*, HanXRQ: *Helianthus annuus*, MD: *Malus domestica*, Mapoly: *Marchantia polymorpha*, Medtr: *Medicago truncatula*, Migut: *Mimulus guttatus*, GSMUA: *Musa acuminata*, OIT: *Nicotiana attenuata*, GWHPAAYW: *Nymphaea colorata*, LOC_Os: *Oryza sativa japonica*, Peaxi: *Petunia axillaris*, Pp: *Physcomitrella patens*, MA: *Picea abies*, Potri: *Populus trichocarpa*, Semoe: *Selaginella moellendorffii*, Seita: *Setaria italica*, Solyc: *Solanum lycopersicum*, PGSC: *Solanum tuberosum*, Sobic: *Sorghum bicolor*, Thecc: *Theobroma cacao*, VIT: *Vitis vinifera*, Zm: *Zea mays*. (Right Top) Expression profiles for genes of the suberin biosynthetic pathway. Dot sizes represent the percentage of cells in which each gene is expressed (% Exp.); and colors indicate the average scaled expression of each gene in each developmental stage group with warmer colors indicating higher expression levels. R.C.: Root cap. Q.C.: Quiescent center. Col: Columella. Procamb: Procambium. (Right bottom) Expression of *S. lycopersicum* genes in the single cell population. The color scale represents log2-normalized corrected UMI counts.

**Figure S7.**
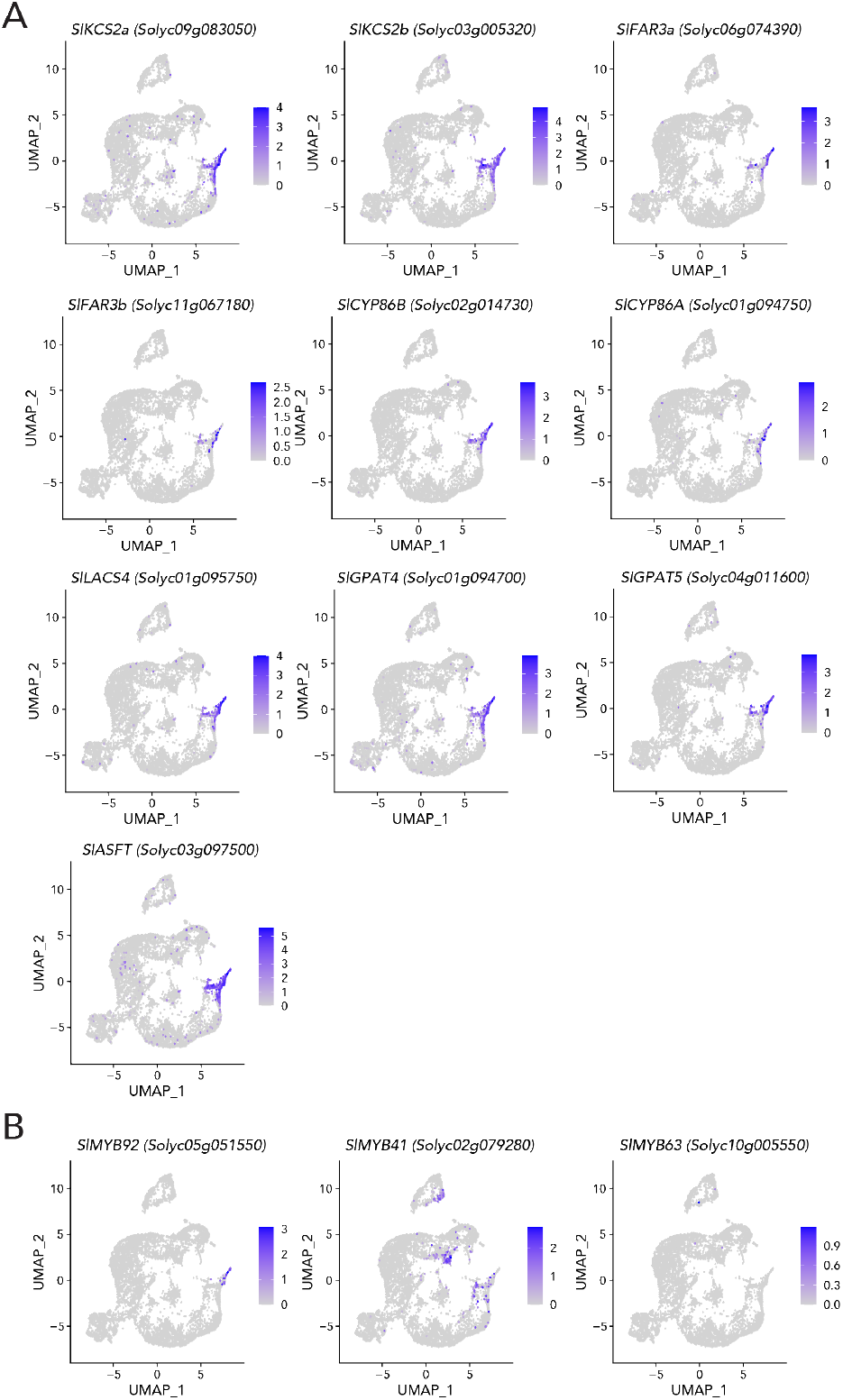
Expression of suberin biosynthetic genes and *SlMYB92* is restricted to the mature exodermis. Expression of candidate (A) biosynthetic genes and (B) transcriptional regulators across the UMAP of Endodermis/Exodermis/Cortex single cell populations. The color scales represent log_2_-normalized corrected UMI counts.

**Figure S8.**
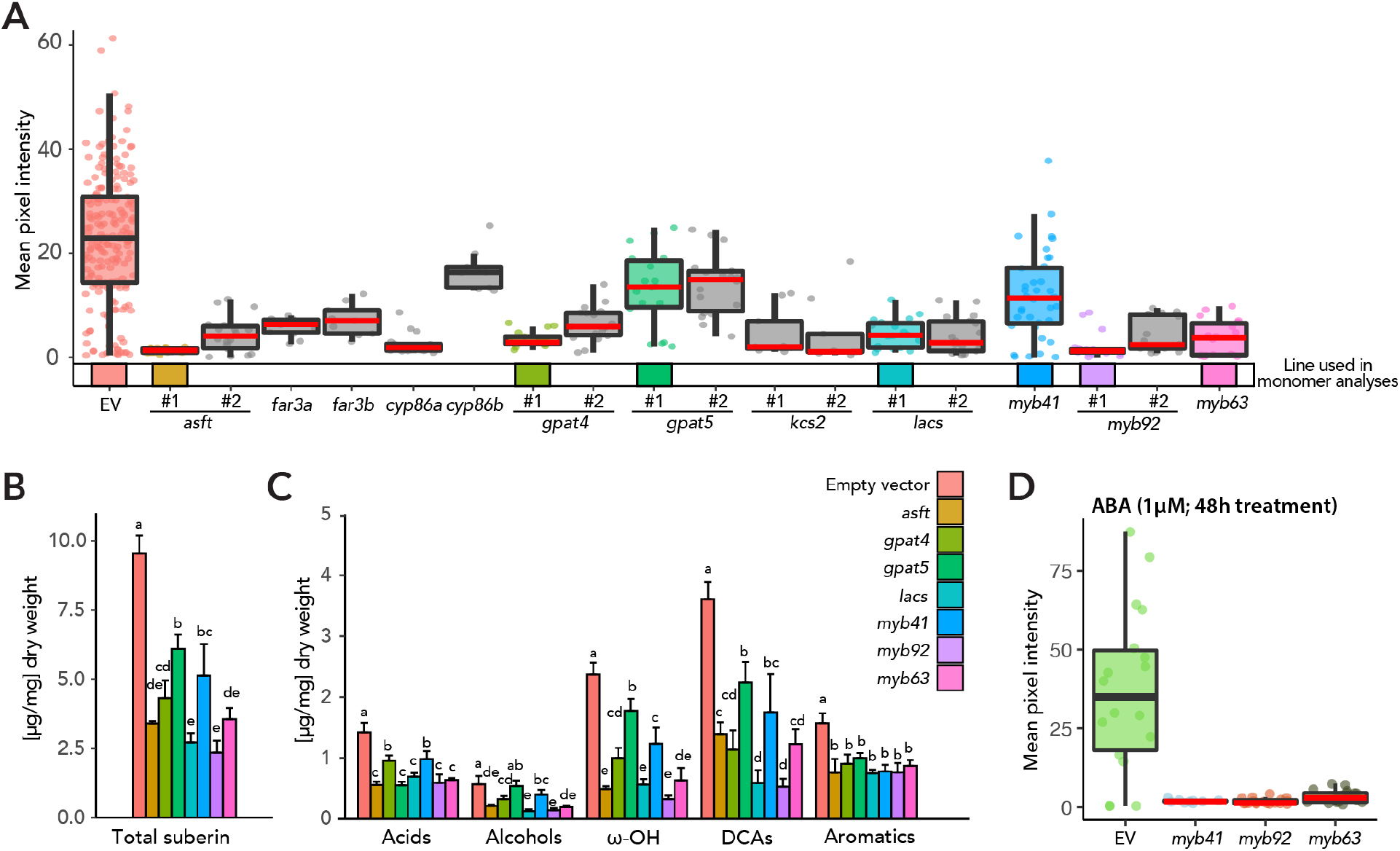
*R. rhizogenes*-derived loss-of-function mutant alleles of candidate genes have impaired suberin deposition. (A) Extended analysis of mutant phenotypes of candidate genes in hairy roots (HR). Quantification of fluorol yellow signal across multiple cross sections (n ≥ 6). Red line indicates statistically significant differences in fluorol yellow pixel intensity in the mutant vs wild type as determined with a one-way ANOVA followed by a Tukey HSD test (adj p-value < 0.05). In most cases, two independently generated HR lines were analyzed, as indicated in the plot. (B) Total suberin abundance and (C) monomer composition of *R. rhizogenes*-generated mutants of suberin biosynthetic enzymes and transcriptional regulators. Acid: fatty acids; Alcohols: primary alcohols; ω-OH: ω-hydroxy fatty acids; DCA: dicarboxylic fatty acid; Aromatics: ferulate and coumarate derivatives. (D) ABA treatment (1 μM for 48 h) does not restore suberin to wild type levels by fluorol yellow staining in *slmyb41*, *slmyb92* and *slmyb63* lines. Mean pixel intensities are not comparable between plots A and D as these were taking under different laser settings.

**Figure S9:**
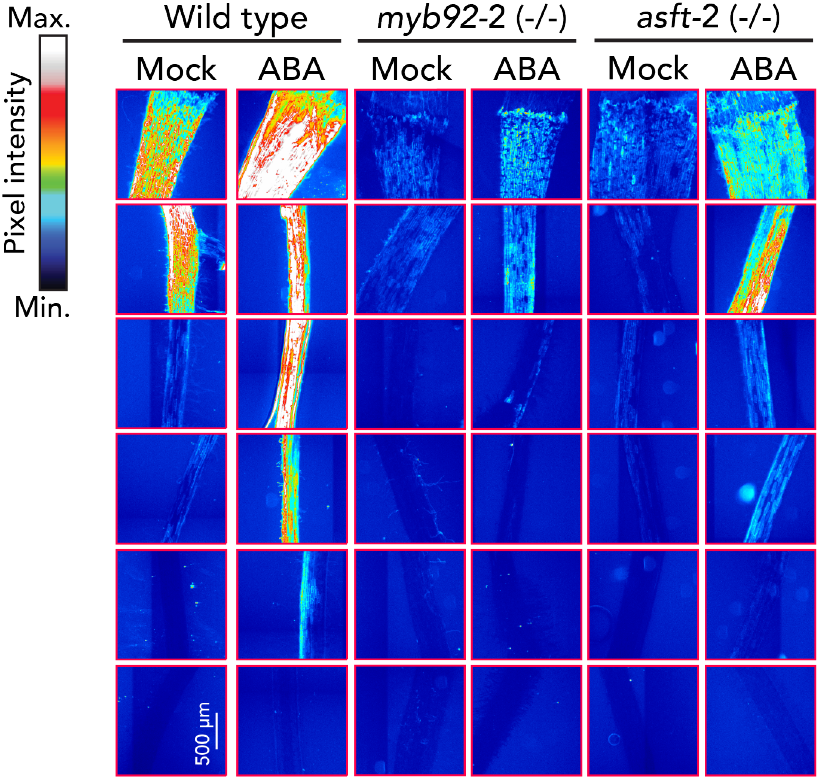
Impaired suberin deposition in the *myb92-2* and *asft-2* alleles and their impact on the response to ABA. Fluorol Yellow (FY) staining for suberin in wild type (repeated from figure 1 for reference), *myb92-2* and *asft-2* plants treated with mock or 1 μM ABA.

**Figure S10:**
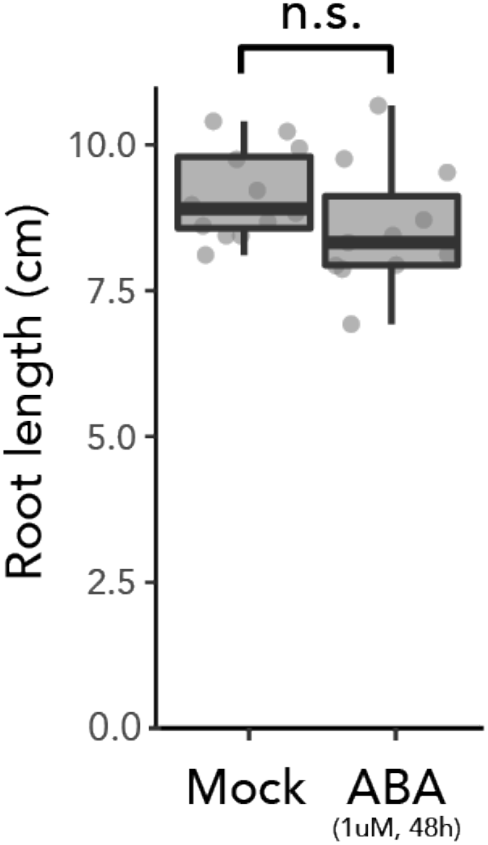
Root length is not significantly affected by the ABA treatment. Boxplot of total root length of 7-day-old wild-type plants treated with mock or 1 μM ABA for 48 h (n=12). A one-way ANOVA analysis did not find any statistically significant differences.

**Figure S11.**
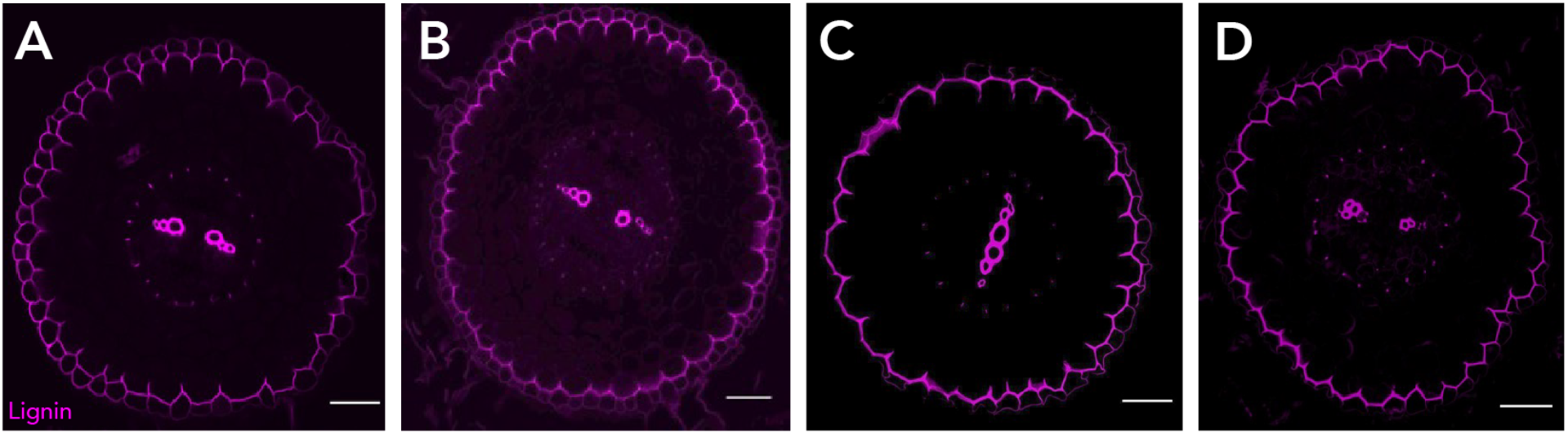
Lignin polar cap in the exodermis is not affected in the *myb41 myb63* and *myb92* hairy root mutants. **A**. Cross section of control hairy root stained with basic fuchsin. **B**. Cross section of *myb92* hairy root mutant stained with basic fuchsin. **C**. Cross section of *myb63* hairy root mutant stained with basic fuchsin. **D**. Cross section of *myb41* hairy root mutant stained with basic fuchsin. Scale bars=50 μm.

**Figure S12:**
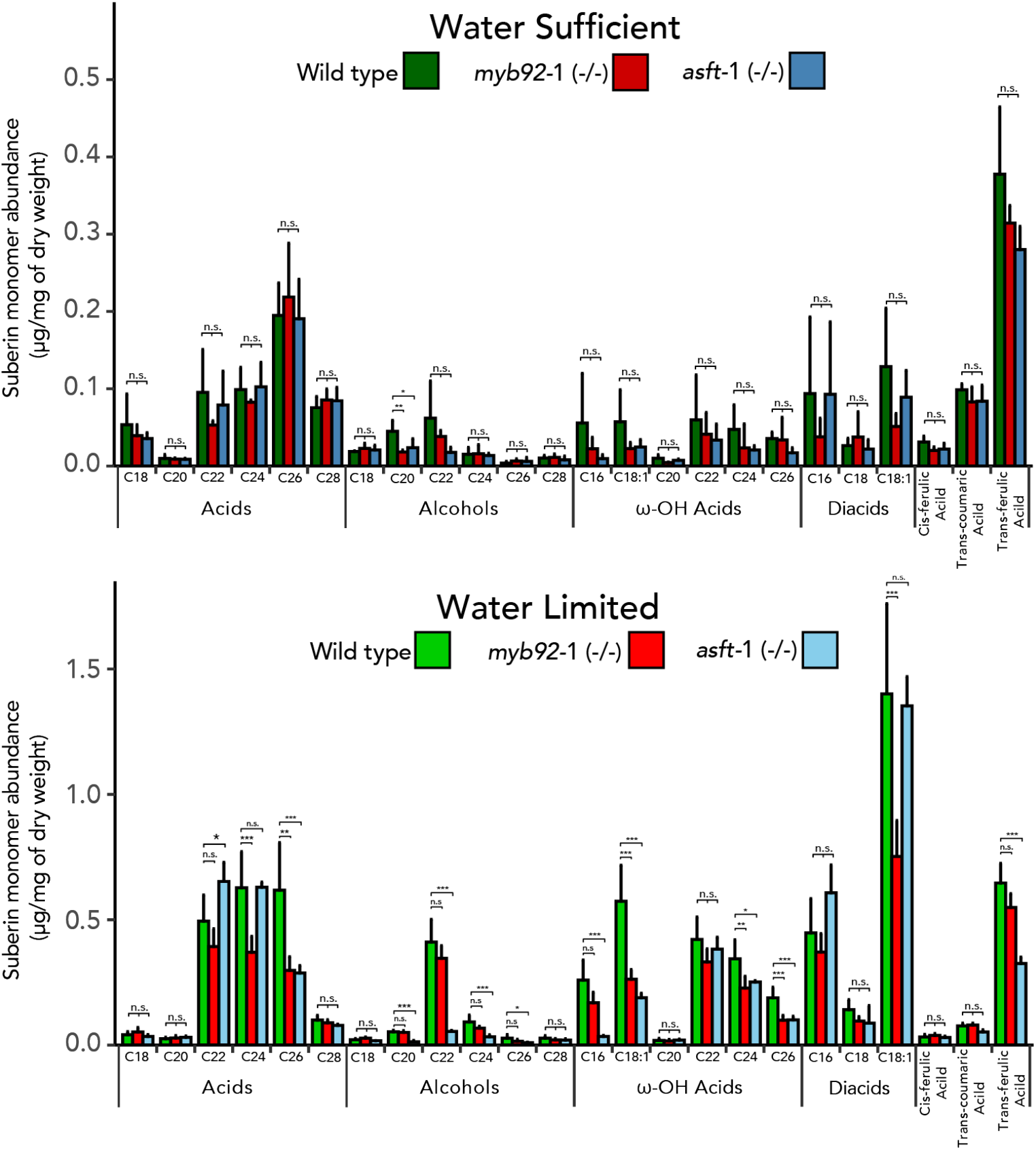
Suberin monomer breakdown of wild type, *myb92-1*, and *asft-1*. Breakdown of specific monomer abundance of samples under water sufficient (top) and water limited conditions (bottom). Acid: fatty acids; Alcohols: primary alcohols; ω-OH: ω-hydroxy fatty acids; DCA: dicarboxylic fatty acid; Aromatics: ferulate and coumarate derivatives. Significance codes: ‘***’ <0.001 ‘**’ <0.01, n.s.: “not significant”.

**Figure S13:**
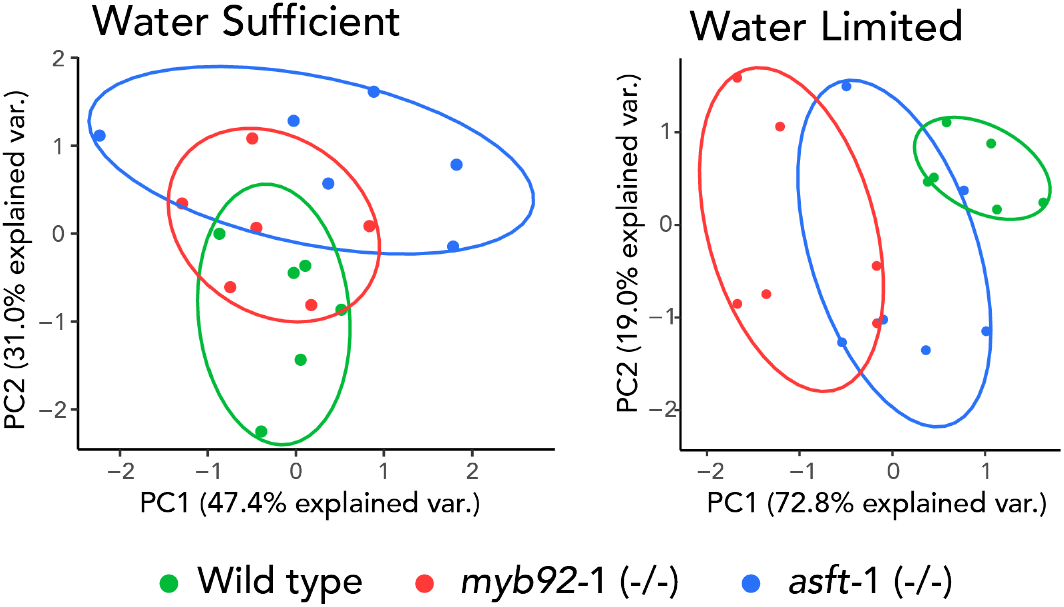
Different overall responses to water limitation in *slasft-1* and *slmyb92-1* compared to wild type. Principal Component (PC) analysis of physiological traits of plants grown in water sufficient, and water limited conditions. Each sample is indicated by a dot and colored by the genotype (n=6).

## SUPPLEMENTARY TABLES

**Table S1**: Canonical suberin biosynthesis enzymes and transcriptional regulators identified in the ‘royal blue’ module. Genes upregulated in fruit that are related to suberin deposition in a DCRi mutant (Lashbrooke et al., 2016).

**Table S2**: Tomato root protoplasting-induced genes.

**Table S3**: Genes activated in the mature exodermal developmental trajectory in the single cell transcriptome data.

**Table S4**: CRISPR design and mutant description. (A) Allele description for *myb92-1, myb92-2, asft-1* and *asft-2*. (B) CRISPR summary of small guide RNA sequences that target PAM sites chosen for this study.

**Table S5**: Oligo sequences used in this study.

## AUTHOR CONTRIBUTIONS

Loosely based on CRediT author statement:

**1.- Conceptualization**: ACP; KK (BSynth candidates); LSM (physiology); SG (initial metabolite); CM (TF candidates and lignin assay); NN (BSynth candidates); DAW (Drought for WGCNA); KAS (physiology); NS (LICOR); JBS (ABA); NG (Substructure); SL (cluster ID); RBF (suberin comp); SMB.

**2.- Methodology**: ACP; PT; SL; RBF; SMB.

**3.- Investigation**: ACP; KK (Figure S4); LSM (Figure 4, S13); CM (Figure S11); DdB (Figure 1, 3, S1); SG (Figure S2); JH (Figure 4, Figure S12); HY (Figure S3, S9, S10); SM (Figure 3, S8); KS (Figure S8); RU (Figure 1, 3, S1); DAW (Figure S4); RBF (Figure 4, S2, S8, S12).

**4.- Computational investigation**: ACP; PT.

**5.- Formal Analysis**: ACP; PT; SL; SMB.

**6.- Resources (Constructs)**: ACP; KK (design Biosynth CRISPR); LSM (design CRISPR); CM (TF HRs); NN (design Biosynth CRISPR); GAM (CRISPR screen); MM (CRISPR screen).

**7.- Writing - original draft**: ACP; KK, DdB; NG; SL; RBF; SMB.

**8.- Visualization/Figures**: ACP; PT; DdB; CM.

**9.- Supervision**: ACP; KK; AB; KAS; NS; NG; SL; RBF; SMB.

**10.- Editing**: All authors contributed to the edition of the manuscript.

## ACKNOWLEDGMENTS

We would like to thank A.M. Bagman and K. Zumstein for experimental assistance; C. Bian, K. Morimoto, and T. Taylor for discussion; D. Kawa and G. Demirer for manuscript critique. Funding was as follows: ACP, SMB, NS, JBS by NSF PGRP IOS-1856749 and IOS-211980. SMB by HHMI 55108506. KK by SKR Postdoctoral Fellowship and MSCA RI Fellowship 790057. LSM by BARD FI-570-2018 and HFSP #RGP0067. CM by MSCA GF 655406. PT and SL by DOE DE-SC0020358 and NSF MCB-2218234. DAW by Elsie J. Stocking Fellowship PBD-UCD. SL by USDA-HATCH program. RU by EMBO ALTF 1046-2015. GAM by NSF PRFB IOS-1907008. ATB and JBS by NSF DGE-1922642. ATB by USDA NIFA 1026477.

## DATA AND RESOURCE AVAILABILITY

Single-cell and bulk RNA-seq data have been deposited at GEO: GSE212405 and will be publicly available at the date of publication. CRISPR-generated mutant lines are available upon request. Timing is dependent upon obtaining phytosanitary certificates according to seed import regulations of the country of destination and associated costs.

